# OMA1-mediated cleavage of AIFM1 upon mitochondrial stress and suppression of cell growth through the control of OXPHOS activity

**DOI:** 10.1101/2025.05.19.654756

**Authors:** Mitsuhiro Nishigori, Serina Hirata, Hidetaka Kosako, Hendrik Nolte, Jan Riemer, Thomas Langer, Takumi Koshiba

## Abstract

Mitochondrial proteases regulate the dynamic properties of organelle morphology and ensure functional plasticity at the cellular level. The metalloprotease OMA1 mediates constitutive and stress-inducible processing of its mitochondrial substrates, but the number of functionally characterized targets remains limited. Using multiproteomic and biochemical approaches, we demonstrated that the membrane-anchored inner membrane space (IMS) protein AIFM1 serves as a mitochondrial stress-responsive substrate of OMA1. OMA1 cleaves AIFM1 in the IMS under stress conditions with a kinetically slower reaction than that of its conventional substrate, dynamin-like GTPase OPA1. Membrane dislocation of cleaved AIFM1 in mitochondria reduces its binding to subunits of the oxidative phosphorylation machinery, leading to decreased respiratory activity and ultimately impaired cell growth. Mechanistically, we revealed that AIFM1 broadly safeguards the mitochondrial proteome under steady-state conditions by mediating the import of proteins, particularly respiratory complex I subunits, via the TIM23 complex. These findings reveal a previously unrecognized role for OMA1 in integrating mitochondrial stress sensing and cellular energetics by altering the AIFM1 topology.

## Introduction

Mitochondria are remarkably dynamic organelles in eukaryotic cells that undergo continuous cycles of homotypic fusion and fission events. Maintaining proper mitochondrial morphology is essential not only for accurate organelle inheritance but also for optimizing physiologic functions, such as signal transduction, metabolic activity, and quality control (Wai & Langer, 2016; Ng *et al*, 2021; Picard & Shirihai, 2022; Tábara *et al*, 2025). Recent studies in an *in vivo* model revealed that genetic ablation of individual components involved in mitochondrial dynamics impairs their organ functions and whole-body metabolism (Chen *et al*, 2003; Ishihara *et al*, 2009; Wakabayashi *et al*, 2009; Wai *et al*, 2015; Tezze *et al*, 2017). Abnormal mitochondrial dynamics are also associated with some human diseases or disorders, such as Charcot-Marie-Tooth disease type 2A (Züchner *et al*, 2004), autosomal dominant optic atrophy (Delettre *et al*, 2000), neuromuscular defects (Shamseldin *et al*, 2012), and a lethal developmental disorder (Waterham *et al*, 2007). In addition to these inherited disorders, an imbalance in mitochondrial fusion and fission can suppress innate immune responses to viral infection (Castanier *et al*, 2010; Koshiba *et al*, 2011; Hanada *et al*, 2020; Yasukawa *et al*, 2020), highlighting the intimate link between mitochondrial dynamics and the host defense system.

Defects in mitochondrial fusion significantly enhance the heterogeneity and dysfunction of the mitochondrial population within the cell (Chen *et al*, 2005), leading to impaired stress responses and loss of cellular homeostasis (MacVicar & Langer, 2016). In mammals, three conserved guanosine triphosphatases (GTPases) of the dynamin family coordinate the fusion process: Mitofusins (Mfn1 and Mfn2) localized to the mitochondrial outer membrane (MOM) and optic atrophy 1 (OPA1), localized to the mitochondrial inner membrane (MIM). In humans, OPA1 has eight isoforms that derive from alternative splicing variants with further post-translational processing by mitochondrial proteases (Wai *et al*, 2015; Chan, 2020). Overlapping with the *m*-AAA protease 1 homolog (OMA1) is a stress-activated MIM peptidase that belongs to the M48 family of zinc metallopeptidases (Käser *et al*, 2003; López-Pelegrín *et al*, 2013) and constitutively processes OPA1 under normal conditions (Ishihara *et al*, 2006; Song *et al*, 2007; Ehses *et al*, 2009; Head *et al*, 2009; Quirós *et al*, 2012). General cellular stresses (e.g., depolarization, oxidization, or heat shock), however, lead to OMA1 activation and complete processing of long-form OPA1 (L-OPA1) isoforms in mitochondria (Ehses *et al*, 2009; Head *et al*, 2009; Quirós *et al*, 2012; Anand *et al*, 2014; Baker *et al*, 2014; Murata *et al*, 2020). Various studies of *OMA1* gene ablation in mice revealed its physiologic role of OMA1-regulated pathways. Knockout of *OMA1* causes relatively mild phenotypes showing increased body weight with defective thermogenesis (Quirós *et al*, 2012), but it shows much more vulnerable in different background mice, such as mice model in mitochondrial cardiomyopathy due to an oxidative phosphorylation (OXPHOS) deficiency (Ahola *et al*, 2022) or simultaneous loss of Parkin leading to imbalance of mitochondrial dynamics (Yamada *et al*, 2025). In contrast, in mouse models of neurodegeneration (Korwitz *et al*, 2016), ischemic kidney injury (Xiao *et al*, 2014), or heart failure (Wei *et al*. 2015; Acin-Perez *et al*. 2018), *OMA1* depletion is protective, supporting the pro-apoptotic function of OMA1 by preventing OPA1 cleavage under stress conditions. Thus, OMA1 manipulates cell survival in a cell- and tissue-specific manner, although the exact mechanism is currently unknown. Recent studies have further implicated OMA1 in another mitochondrial signaling pathway that activates the integrated stress response via DAP3-binding cell death enhancer 1 (DELE1), a protein kinase activator (Fessler *et al*, 2020 & 2022; Guo *et al*, 2020). In this pathway, stress-activated OMA1 cleaves DELE1 as a substrate during import, generating a short form (S-DELE1) that is released from the mitochondria into the cytosol, where the S-DELE1 ultimately triggers the ISR. Therefore, OMA1 regulates the dynamic properties of mitochondrial morphology and cellular stress signaling to ensure mitochondrial integrity and cellular homeostasis. Despite its emerging role as a central pivotal protease involved in a wide range of mitochondrial homeostatic processes, only a limited number of OMA1 substrates have been functionally characterized.

In the present study, using a proteome-wide approach combined with biochemical analysis, we identified the intermembrane space (IMS) protein apoptosis-inducing factor mitochondria associated 1 (AIFM1) as a previously unidentified OMA1 substrate under mitochondrial stress. We demonstrate that OMA1-mediated processing of AIFM1 induces the release of AIFM1 from the MIM and reduces its binding to OXPHOS machinery subunits, ultimately impairing respiration and cell growth. These findings reveal a previously unrecognized role for OMA1 in integrating mitochondrial stress sensing and cellular energetics by altering the topology of one of its substrates.

## Results

### Screening of the OMA1 mitochondrial interactome by mass spectrometry

Because OMA1 is involved in a wide range of biologic processes, from quality control to cellular stress responses (Rivera-Mejías *et al*, 2023), we reasoned that unidentified molecules in the mitochondria might functionally and physically associate with the peptidase to control mitochondrial integrity. To test this hypothesis, we sought to identify proteins that interact with OMA1 as candidate substrates. We first established cell lines stably expressing a carboxyl-terminal Myc-tagged murine OMA1 (termed OMA1/Myc) in *OMA1*-null mouse embryonic fibroblasts (MEFs), validated the properties of the recombinant protein (Fig. EV1A-I), and screened for molecules that interact with OMA1/Myc in cells.

To identify OMA1-associated proteins, we isolated mitochondria from the OMA1/Myc-expressing cells, and after moderate cross-linking with formaldehyde, immunoprecipitated (IP) the OMA1/Myc with an anti-Myc antibody. Liquid chromatography-tandem mass spectrometry (LC-MS/MS) analysis of the precipitated proteins identified 38 proteins that were enriched in the OMA1/Myc-expressing cell immunoprecipitate (Fig. 1A and Dataset EV1), including the known OMA1 substrate and interactor OPA1, the prohibitin membrane scaffold complexes PHB1 and PHB2, and components of the mitochondrial contact site and cristae organizing system (MICOS; MIC60, MIC19, MIC25, and MIC27). Other IMS proteins, HTRA2/Omi and mitochondrial adenylate kinase 2 (AK2), both associated with mitochondrial stress signaling (Chan, 2005; Lee *et al*, 2007), were also enriched in the OMA1 precipitates. In addition to these interactors, AIFM1 was highly enriched (> 16-fold change) (Fig. 1A, red), and this was further validated using the parallel reaction monitoring (PRM) method (Fig. 1B). These results suggest that several molecules, including AIFM1, are members of the OMA1 interactome in mitochondria and are thus candidate substrates of the peptidase.

**Figure 1.**
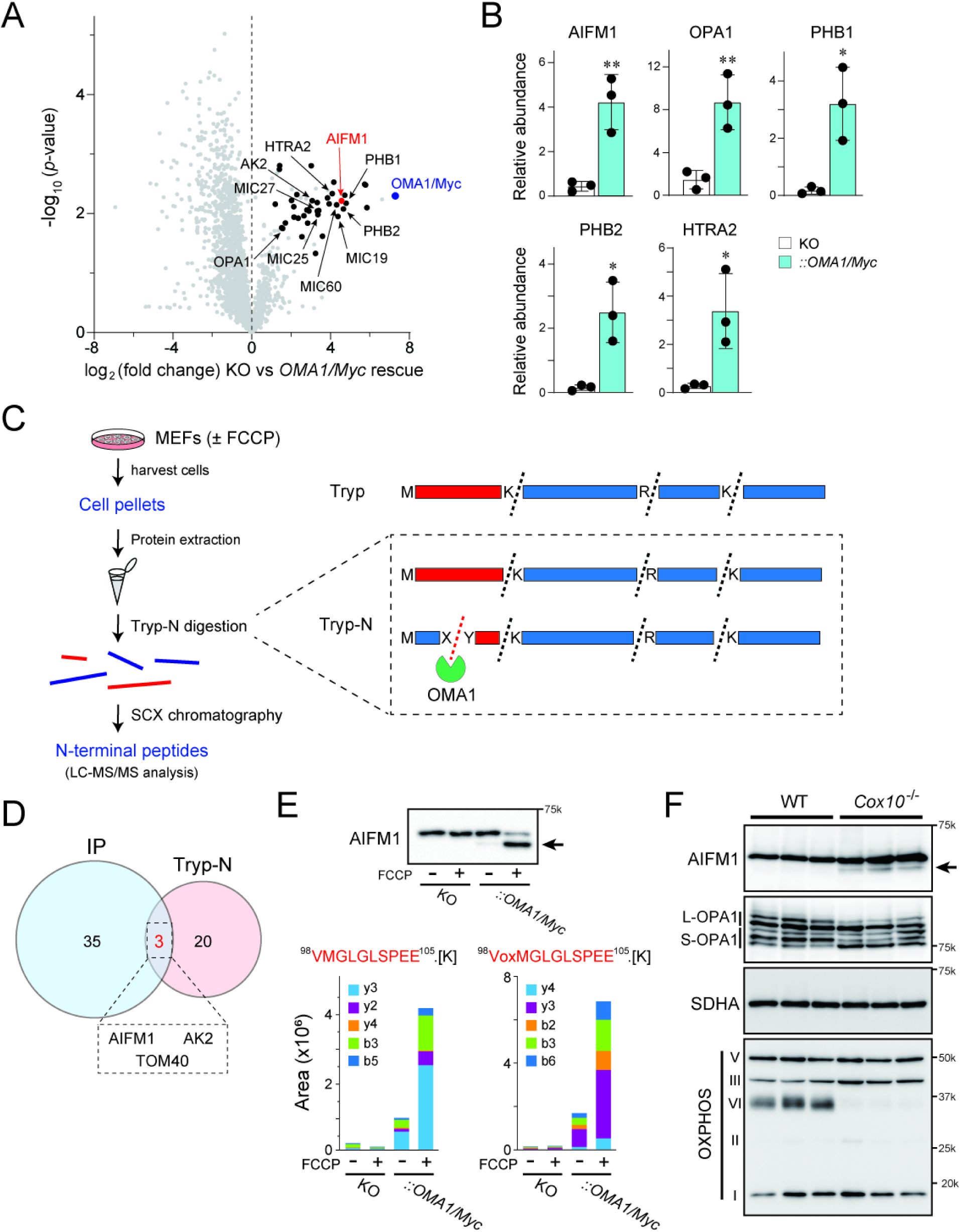
AIFM1 is a stress-inducible OMA1 substrate. **A** Volcano plot showing OMA1/Myc interacting partners. Mitochondrial extracts from *OMA1* KO cells with or without OMA1/Myc expression were immunoprecipitated. Co-purifying proteins were identified by quantitative mass spectrometry (MS) (*n* = 3 biologic replicates). Enriched mitochondrial proteins (abundance ratio >2, *p* < 0.05) are plotted in black showing AIFM1 (red) and OMA1/Myc (blue). Several known IMS or IMM proteins are also indicated. See also Dataset EV1. **B** Relative abundance of proteins (AIFM1, OPA-1, PHB1, PHB2, and HTRA2) used for proteomics in (A) was measured by targeted LC-MS/MS using the PRM method. Data shown are mean values ± S.D. (*n* = 3 biologic replicates). *, *p* < 0.05; **, *p* < 0.01. **C** Schematic of the N-terminal proteomic approach. Protein extracts from MEFs treated with or without FCCP were digested with Tryp-N peptidase, followed by separation of the reactants by strong cation exchange (SCX)-based chromatography, and analysis of the purified peptide fragments by LC-MS/MS. In this system, each Tryp-N-digested peptide fragment would have at least one lysine or arginine residue at the N-terminus (right inset), but that of the OMA1 substrates does not follow this rule (recovered in the flow-through in SCX chromatography). **D** Venn diagram showing the overlap (3 candidates; AIFM1, AK2, and TOM40) between the indicated mitochondrial molecules revealed by IP-MS (38 candidates) and the N-terminal proteomic approach (23 candidates). See also Table EV1 and Dataset EV2. **E** The Tryp-N digested AIFM1 peptide (^98^VMGLGLSPEE^105^) from the *OMA1* KO or the *OMA1/Myc* rescued MEFs with or without FCCP treatment was quantified by targeted MS using the PRM method (the samples used in [C]). Note that the peptide was enriched only in the rescued sample and lacked K or R residues at the N-terminus, indicating that the fragment generation was OMA1-dependent. oxM, oxidized methionine. Top blot shows the immunoblot of the samples used in the study (detected by anti-AIFM1 antibody). The arrow indicates a generated AIFM1 cleavage product. **F** Loss of Cox10 activates OMA1 and leads to increased AIFM1 processing *in vivo*. Heart lysates from WT or *Cox10*^−/−^ mice (*n* = 3) were analyzed by immunoblotting (indicated antibodies). The arrow indicates the AIFM1 processed band.

### Mitochondrial stress triggers OMA1-dependent processing of AIFM1

To elucidate unidentified substrates of OMA1 during mitochondrial stress, we then used a proteome-wide approach to identify neo-amino-terminal peptides that are formed in an OMA1-dependent manner. MEFs expressing OMA1/Myc or *OMA1* knockout (KO) cells treated with or without carbonyl cyanide-4-(trifluoromethoxy)phenylhydrazone (FCCP) were lysed, and their total protein extracts were digested with the thermostable protease LysargiNase (Tryp-N; Zhou *et al*, 2019), followed by proteome analysis (Fig. 1C). We observed peptide enrichment that was specific for a stress response (± FCCP) coupled with an OMA1 dependency (KO vs *OMA1/Myc* rescue) and identified 23 unique peptides derived from total mitochondrial proteins (Table EV1 and Dataset EV2). Overlaying this peptide enrichment profile with the previous immunoprecipitation-mass spectrometry (IP-MS) result (Fig. 1A), we identified three mitochondrial proteins (AIFM1, AK2, and TOM40) as OMA1 substrate candidates under mitochondrial depolarization (Fig. 1D).

Among these candidates, substantial AIFM1 processing in response to FCCP treatment in *OMA1/Myc*-rescued cells was confirmed by immunoblotting (Fig. 1E, arrow in top blot), consistent with data quantified by targeted MS using the PRM method (bottom graphs). In contrast to OMA1/Myc, expression of the OMA1^E324Q^ mutant in *OMA1* KO cells did not produce AIFM1 processing under depolarized conditions (Fig. EV2A). Hyperpolarization induced by oligomycin A (ATP synthase inhibitor) also triggered AIFM1 proteolysis in an OMA1-dependent manner (Fig. EV2B). These results demonstrate OMA1-dependent processing of AIFM1 under stress conditions. To corroborate these *in vitro* findings, we analyzed AIFM1 processing in tissue from the hearts of *Cox10*^−/−^ mice. Loss of Cox10 is associated with cardiomyopathy (Diaz *et al*, 2005) and leads to the activation of OMA1 and increased OPA1 processing (Ahola *et al*, 2022; Fig. 1F); importantly, we observed at least some AIFM1 processing in cardiac tissue (Fig. 1F, arrow).

Several mitochondrial proteases process their substrates under constitutive and/or stress-induced conditions. The IMS-localized *i*-AAA protease YME1L and OMA1 share substrates, such as L-OPA1, although these proteases recognize distinct sites in the protein (Deshwal *et al*, 2020). In terms of redundancy, we investigated whether YME1L might also proteolyze AIFM1 under certain circumstances. By monitoring AIFM1 proteolysis in both *YME1L*^−/−^ and *OMA1*^−/−^*YME1L*^−/−^ (DKO) cells under constitutive (DMSO) or depolarized (FCCP or valinomycin) conditions, we confirmed that the AIFM1 cleaved band was only observed in cells with OMA1 (Fig. 2A, wild-type [WT] and *YME1L*^−/−^). Consistent with this finding, reintroduction of *OMA1*, but not *YME1L*, into the DKO cells was sufficient to trigger AIFM1 proteolysis (Fig. EV2C), suggesting that these two proteases are functionally distinct, with respect to AIFM1 cleavage. We further investigated the actions of other mitochondrial proteases that may be involved in AIFM1 proteolysis. Knockdown of five independent IMS or MIM proteases in HeLa cells by small interfering RNA (siRNA) strongly attenuated the level of each protein (Fig. 2B). Although the FCCP-induced AIFM1 cleavage was clearly inhibited in *siOMA1*-treated cells, as observed above in *OMA1*^−/−^ cells, knockdown of the other four proteases (YME1L, AFG3L2, HTRA2, and PARL) had no effect (Fig. 2B, top). Thus, we concluded that AIFM1 is a substantial OMA1 substrate under mitochondrial stress conditions.

**Figure 2.**
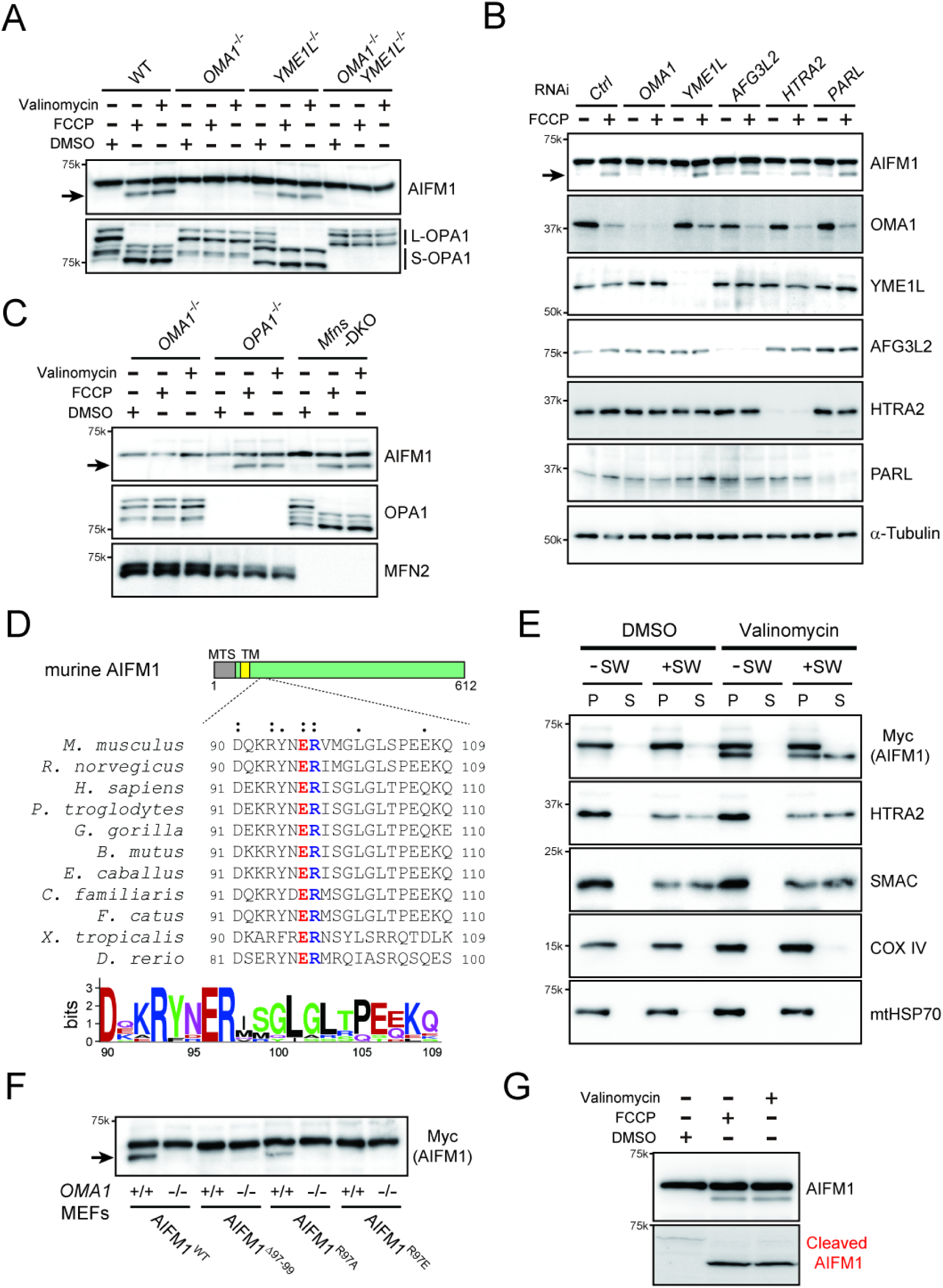
OMA1-dependent processing of AIFM1 in cells. **A** OMA1 is required for inducible AIFM1 processing. WT, *OMA1*^−/−^, *YME1L*^−/−^, or *OMA1*^−/−^*YME1L*^−/−^ MEFs were incubated for 3 h in the absence (DMSO) or presence of either FCCP (40 μM) or valinomycin (1 μg/mL) and analyzed by immunoblotting (indicated antibodies). The arrow indicates the AIFM1 processed band. **B** HeLa cells were transfected with the indicated siRNA and treated with or without FCCP (40 μM) for 3 h. Immunoblotting was performed as indicated and the arrow indicates the AIFM1 processed band. **C** Similar to (A), except that *OPA1*^−/−^ and *Mfn1*^−/−^*Mfn2*^−/−^ (*Mfns*-DKO) MEFs were used. *Mfns*- DKO and *OPA1*-KO are incompetent for OM and IM fusion, respectively. **D** Alignment of AIFM1 homologs (*Mus musculus*, *Rattus norvegicus*, *Homo sapiens*, *Pan troglodytes*, *Genus gorilla*, *Bos mutus*, *Equus caballus*, *Canis familiaris*, *Felis catus*, *Xenopus tropicalis*, and *Danio rerio*) downstream of the transmembrane (TM) region. Colons (:) indicate conserved residues among all species shown in the figure, and dots (·) indicate residues with similar properties. Top, structure of murine AIFM1 showing the location of the mitochondrial target signal (MTS) and the TM segment, with amino acid positions indicated below the structure. Bottom, sequence logo of positions adjacent to the cleavage site in AIFM1 by OMA1. **E** Mitochondria isolated from WT MEFs treated with or without valinomycin (1 μg/mL) were diluted in either isotonic (−SW) or hypotonic swelling (+SW) buffer and kept on ice for 15 min. After centrifugation, the supernatant (S) and pellet (P) were analyzed by Western blotting using the indicated sub-mitochondrial markers. HTRA2 and SMAC are used as a positive control for unanchored IMS protein. COX IV and mtHSP70 are MIM and matrix proteins, respectively. **F** MEFs (*OMA1*^+/+^ or *OMA1*^−/−^) stably expressing a Myc-labeled version of WT or mutant (Δ97-99, R97A, or R97E) AIFM1 were treated with 40 μM FCCP for 3 h and analyzed by immunoblotting with anti-Myc antibody. The arrow indicates the AIFM1 cleaved band. **G** The cleaved AIFM1 antibody (bottom) specifically recognizes proteins with the identified cleaved end of the N-terminus. The WT MEF samples used in (A) were tested.

### Mechanistic study of OMA1-mediated AIFM1 processing

As the functional domains of both OMA1 and AIFM1 are directed toward the IMS and are anchored to the MIM, we next investigated how OMA1 mechanistically accesses its substrate during mitochondrial proteolysis. To address this issue, we first examined the dynamic properties of the mitochondrial membrane using fusion- or fission-incompetent cells. Interestingly, mitochondrial depolarization induced by either FCCP or valinomycin in *OPA1-* or *Mitofusins* (*Mfn1*/*Mfn2*)-deficient cells was sufficient to trigger the OMA1-dependent processing of AIFM1 (Fig. 2C). In addition, *Drp-1*^−/−^ MEFs, which lack the mitochondrial fission event (Ishihara *et al*, 2009; Wakabayashi *et al*, 2009), also exhibited normal AIFM1 processing through OMA1 function (Fig. EV2D). These results suggest that recognition of AIFM1 by OMA1 is likely to be dispensable for both mitochondrial fusion and fission processes (Fig. EV2E).

Our peptide mapping based on the N-terminal proteome identified that the valine at position 98 was the N-terminal amino acid of the generated AIFM1 fragment cleaved by OMA1 (Fig. 1E and Table EV1), and the cleavage site was located just downstream of the transmembrane domain in AIFM1 (Fig. 2D, top structure, and Fig. EV3A). Consistent with this observation, cleaved AIFM1 was released from the MIM into the IMS, where it existed as a soluble, membrane-unbound protein like HTRA2/Omi or SMAC/DIABLO (Fig. 2E). Sequence alignments between

AIFM1 homologs in different species revealed conservation of the OMA1 recognition sequences, particularly the Glu^96^-Arg^97^ dipeptide just before the cleavage site (Fig. 2D), where we observed no conservation of any OMA1 cleavage motif (Fig. EV3B) (Ishihara *et al*, 2006; Guo *et al*, 2020; Ahola *et al*, 2024). We generated an AIFM1 deletion mutant lacking a tripeptide motif that includes Val^98^ (AIFM1^Δ97-99^, Fig. EV3C), and tested whether OMA1 could properly cleave the mutant protein. As expected, OMA1 failed to cleave the deletion mutant in cells (Fig. 2F and EV3D). We also introduced substitutions into AIFM1 at the site corresponding to the OMA1 recognition sequence and found that the R97E variant could not be processed by OMA1 (Fig. 2F and EV3D). These results demonstrate that the tripeptide motif in AIFM1 is essential for its recognition and cleavage by OMA1, and that the net charge surrounding the upstream cleavage site in AIFM1 likely facilitates substrate recognition.

Finally, we generated a specific antibody against the cleaved form of AIFM1 to validate our biochemical findings of the cleavage site at the endogenous level (Fig. EV4A-B). Using this antibody, we detected a cleaved AIFM1 band with an apparent molecular mass of approximately 56 kDa, but only under conditions in which the mitochondrial membrane potential (ΔΨ_m_) was dissipated (Fig. 2G, bottom). Notably, the antibody-based probe was also highly sensitive to a range of mitochondrial stresses known to activate OMA1 (Fig. EV4C-D).

### Differences in the rates of OMA1-mediated cleavage of AIFM1 and OPA1

When WT MEFs were treated with FCCP, activated OMA1 rapidly attacked L-OPA1 and induced degradation of the substrate within a few minutes, leading to the accumulation of S-OPA1 isoforms, while the OPA1 pattern in *OMA1*^−/−^ cells was not altered (Fig. 3A, middle blots) (Baker *et al*, 2014). In contrast, OMA1-mediated processing of AIFM1 occurred significantly more slowly than that of OPA1, becoming apparent only approximately 15 min after membrane depolarization (Fig. 3A-B). It is conceivable that OMA1 only cleaves AIFM1, whose import into depolarized mitochondria is inhibited (Chacinska *et al*, 2009). Inhibition of cytosolic protein synthesis with cycloheximide, however, did not alter the kinetics of OMA1-mediated processing of AIFM1 or OPA1 (Fig. 3C, CHX), excluding the possibility that the observed kinetic difference between these substrates was due to the import failure of the substrates upon mitochondrial depolarization.

**Figure 3.**
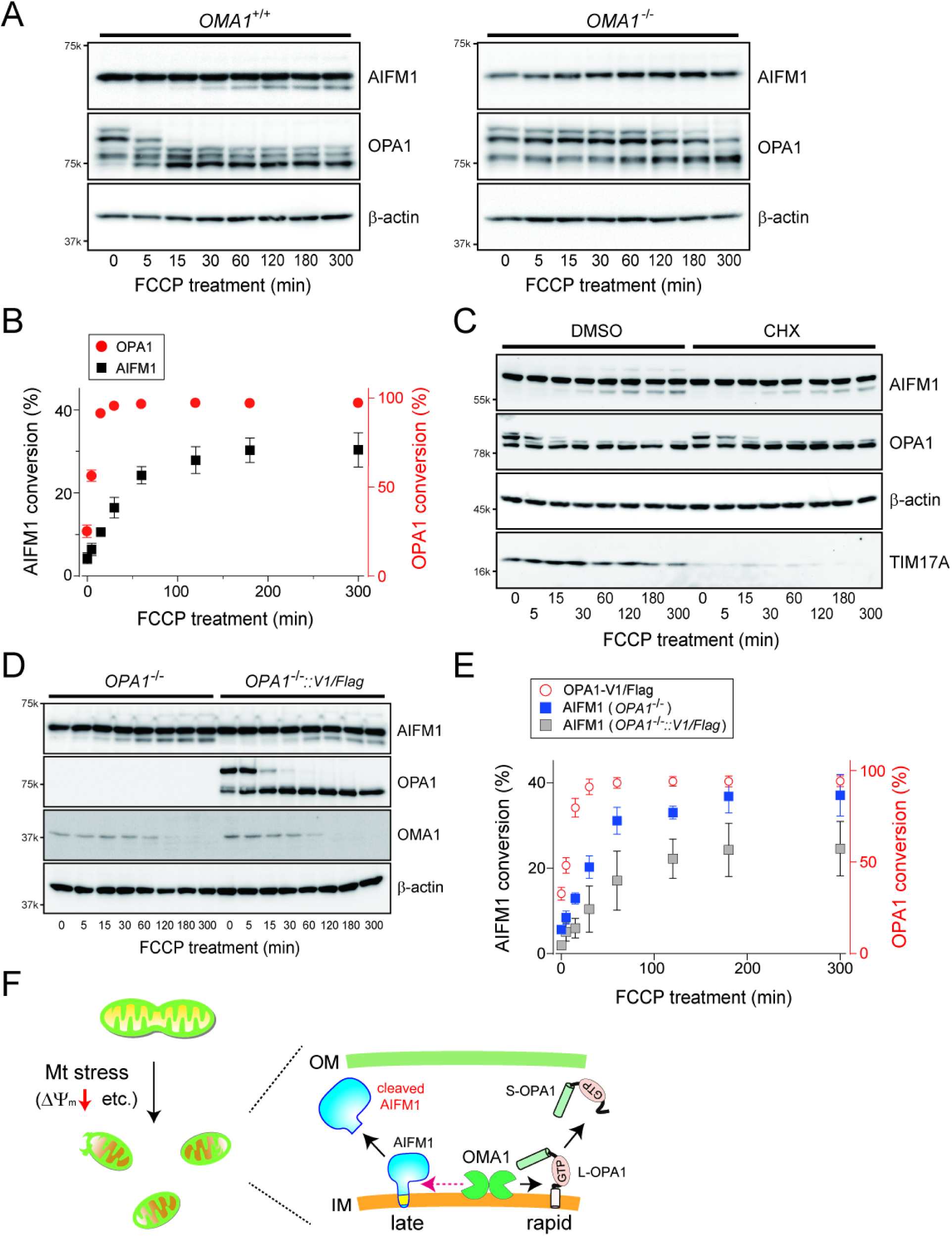
AIFM1 processing by OMA1 is a relatively slow reaction. **A** The kinetic profile of stress-induced substrate processing (OPA1 and AIFM1) is dependent on OMA1. The *OMA1*^+/+^ (left) or *OMA1*^−/−^ (right) MEFs incubated with FCCP (40 μM) were collected at the indicated time points (0, 5, 15, 30, 60, 120, 180, and 300 min) and analyzed by Western blotting. β-Actin blots were used as loading controls for each time point. **B** OMA1-dependent substrate processing in *OMA1*^+/+^ MEFs (A) was quantified (*n* = 3 biologic replicates) and conversions (%) were plotted. **C** Similar to (A), except that *OMA1*^+/+^ MEFs pretreated with either DMSO or cycloheximide (CHX, 10 μg/mL) were used for the FCCP-inducible substrate processing assay. The TIM17A blot was used as a marker to verify the effect of CHX in cells. **D** Similar to (A), except that the *OPA1*^−/−^ and *OPA1*^−/−^ MEFs stably expressing OPA1_variant 1/FLAG were treated with FCCP for the indicated times. **E** The OMA1-dependent AIFM1 processing in the *OPA1*^−/−^ (blue) or *OPA1*^−/−^ MEFs stably expressing OPA1_V1/FLAG (gray) shown in (D) was quantified (*n* = 3 biologic replicates) and each conversion (%) was plotted. The kinetic profile of OPA1_V1/FLAG (red circle) shows the OMA1-dependent processing of the protein in the rescue cells. **F** Model of the AIFM1 processing by OMA1 in the IMS. Mitochondrial stress-induced activation of OMA1 triggers rapid processing of its substrate OPA1 followed by subsequent targeting of AIFM1.

We next asked whether other OMA1 substrates might influence the relatively slow kinetics of AIFM1 processing by OMA1. We removed OPA1, the rapidly processed OMA1 substrate, from the mitochondrial membrane and examined whether the kinetics of AIFM1 processing were altered. Strikingly, AIFM1 cleavage upon mitochondrial depolarization was accelerated in *OPA1*-deficient cells (Fig. 3D). This effect was suppressed in cells expressing either L-OPA1 (variant 1 isoform, Fig. 3D-E) or a non-cleavable OPA1 variant (OPA1V1Δ4, Fig. EV5A) (Ahola *et al*, 2024), demonstrating that S-OPA1 is dispensable for accelerated AIFM1 processing in depolarized mitochondria. In addition, AIFM1 showed an increased susceptibility to OMA1 in the *OPA1*^−/−^ cells, and the protein was degraded at even low levels of mitochondrial stress (FCCP < 1 μM, Fig. EV5B). These observations in the *OPA1*^−/−^ cells were not simply due to cristae morphology defects (Pernas & Scorrano, 2016) or the lose of respiratory activity (Chen *et al*, 2005), as the elimination of MIC60 a core component of MICOS (van der Laan *et al*, 2016) still resulted in a distinct, relatively slower AIFM1 processing kinetic pattern than that in the *OPA1* KO phenotype (Fig. EV5C). Elimination of another OMA1 substrate, DELE1, also had no effect on the kinetics of OMA1-mediated AIFM1 cleavage (Fig. EV5D), suggesting that mitochondrial stress-induced substrate processing events by the activated OMA1 are independent (Fig. 3F).

### Membrane dislocation of AIFM1 affects its complex formation in mitochondria

To further understand the functional relevance of AIFM1 processing, we generated cell lines that accumulate cleaved AIFM1 without inducing mitochondrial stress in the cells. We swapped the arginine 97 position in a carboxyl-terminal Myc-tagged AIFM1 with a tobacco etch virus (TEV) protease cleavage site (TCS; termed AIFM1^TCS^; Fig. 4A) and exploited the Flp-In system to stably express the variant at physiologic levels in the *AIFM1* KO cell lines (Salscheider *et al*, 2022). Insertion of the TCS into AIFM1 did not affect its subcellular localization (Fig. EV6A), and co-expression of TEV protease in the IMS (IMS-TEV) but not the mitochondrial matrix (Su9-TEV) allowed specific processing of AIFM1^TCS^ (Fig. 4B, arrow). Using this system, we produced a mitochondrial protein (designated AIFM1^TCS/TEV^; Fig. 4C-D) with the expected molecular weight (∼60k Da; including Myc tag) at high efficiency (Fig. EV6B). We also confirmed that expression of the TEV protease in the IMS had no significant off-target effects on other mitochondrial proteins, such as OMA1 and OPA1, and did not activate the transcription factor ATF4, which coordinates the mitochondrial stress response in mammals (Fig. EV6C) (Quirós *et al*, 2017; Guo *et al*, 2020; Girardin *et al*, 2021). Having clarified the solubility of the naturally cleaved form of AIFM1 in depolarized mitochondria (Fig. 2E), we found that the AIFM1^TCS/TEV^ variant was similarly released from the MIM, as assessed by the mitochondrial swelling assay (Fig. 4E).

**Figure 4.**
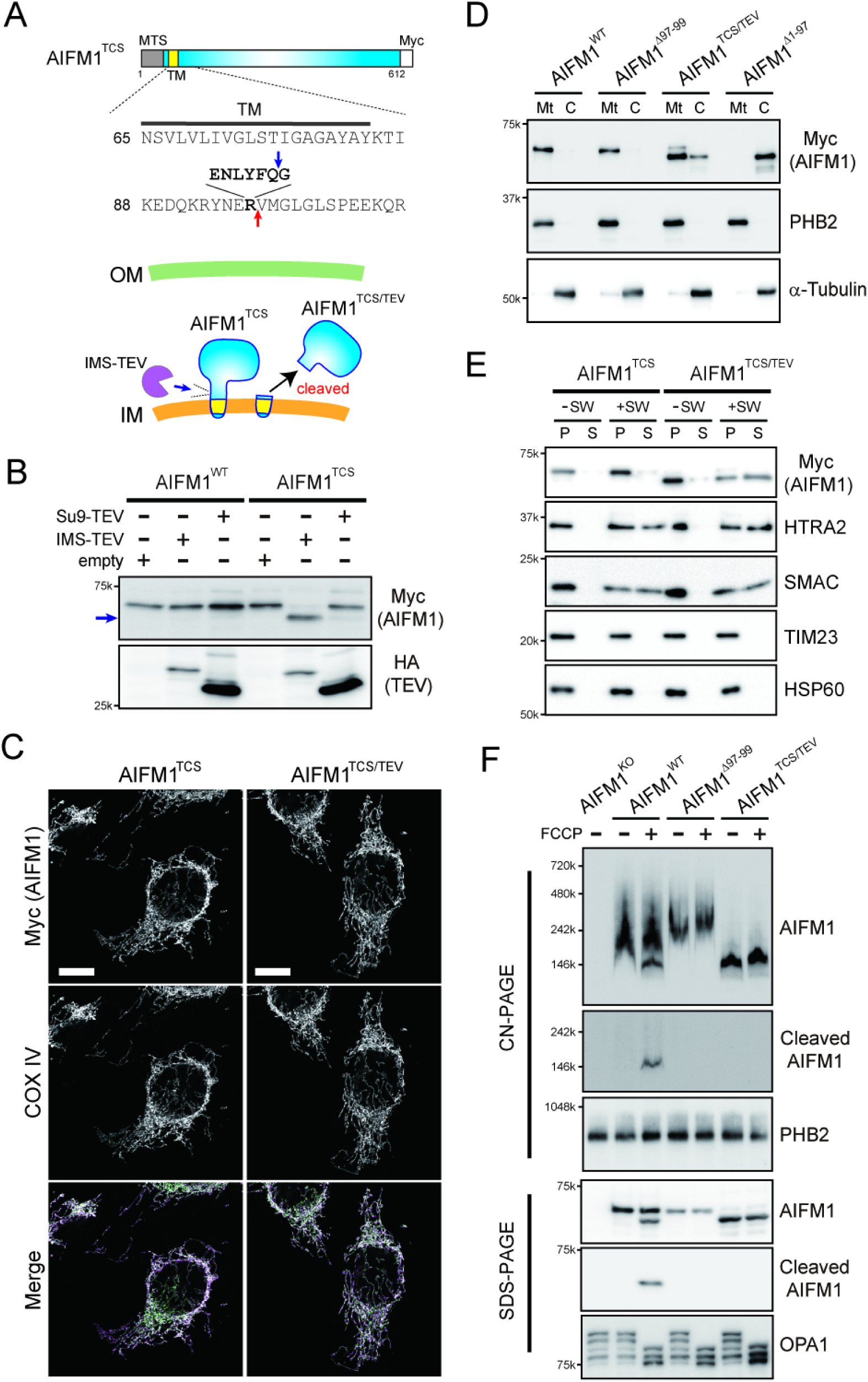
Submitochondrial localization of AIFM1 is necessary to preserve its assembly states. **A** The sequence of AIFM1^TCS^ harboring a TEV cleavage site (TCS) where the R97 residue in AIFM1 has been replaced by TCS (bold). The red arrow in the sequence indicates the original cleavage site by OMA1 and the blue arrow represents the newly generated processing site by the TEV protease. The bottom image illustrates the possible topology of AIFM1^TCS^ in mitochondria and the TEV protease located in the IMS (IMS-TEV) hypothetically cleaving and releasing the variant from the membrane (designated AIFM1^TCS/TEV^). **B** The Flp-In-293-AIFM1^WT^/Myc or - AIFM1^TCS^/Myc cells were transfected with expression plasmids of HA-tagged Su9-TEV (matrix-targeted) or IMS-TEV, and their whole cell lysates were analyzed by Western blotting using the indicated antibodies. The arrow indicates the generated AIFM1^TCS/TEV^. **C** Subcellular localization of AIFM1 variants. Flp-In-293-AIFM1^TCS^/Myc cells without (AIFM1^TCS^) or with stable expression of IMS-TEV using a retroviral system (AIFM1^TCS/TEV^) and the generated AIFM1 variants in the cells were monitored by an immunofluorescence against the Myc epitope to determine their subcellular localizations (top panel). Mitochondria in the same cells were also identified by staining with an anti-COX IV antibody (middle). We confirmed that both AIFM1 (magenta) and COX IV (green) were completely merged in mitochondria (bottom). Scale bar, 10 μm. **D** Cellular fractions from cells expressing the AIFM1 variants (WT, Δ97-99, TCS/TEV, and Δ1-97) were collected by differential centrifugation and analyzed by Western blotting using the indicated subcellular markers (PHB2, mitochondria; α-Tubulin, cytosol). Mt, mitochondrial fraction; C, cytosolic fraction. **E** Similar to Fig. 2E, except that mitochondria were isolated from cells expressing the AIFM1 variants (AIFM1^TCS^ and AIFM1^TCS/TEV^). HTRA2 and SMAC are used as a positive control for an unanchored IMS protein. TIM23 and HSP60 are MIM and matrix proteins, respectively. **F** Depolarized (+ FCCP) or normal mitochondria (− FCCP) isolated from *AIFM1* KO or cells expressing AIFM1 variants (WT, Δ97-99, and TCS/TEV) were solubilized in CN-PAGE lysis buffer containing 0.5% (w/v) digitonin, analyzed by CN-PAGE, and immunoblotted using the AIFM1-, cleaved AIFM1-, and PHB2-specific antibodies (top 3 blots). The abundance of mitochondrial proteins in each sample was also confirmed by SDS-PAGE in parallel (bottom 3 blots).

We found that membrane dislocation of AIFM1 affects its assembly in mitochondria. The soluble AIFM1^TCS/TEV^ variant formed 150-kDa complexes, estimated to be a dimer formation (Maté *et al*, 2002; Ferreira *et al*, 2014) based on clear native (CN)-PAGE (Fig. 4F, right lanes). In contrast, both the membrane-anchored AIFM1^WT^ and non-cleavable AIFM1^Δ97-99^ variants were part of larger complexes with a molecular weight >240 kDa (Hevler *et al*, 2021; Salscheider *et al*, 2022), likely due to the formation of multiple homotypic and/or heterotypic complexes in mitochondria (Fig. 4F). Notably, when the AIFM1^WT^ cells were treated with FCCP (lane, AIFM1^WT^ + FCCP), we observed that two major populations that co-migrated; one with the original larger heterogeneous peaks and the other with a lower mass peak consistent with the predominant band seen in the AIFM1^TCS/TEV^ fraction. Strikingly, our specific antibody to cleaved AIFM1 revealed that the smaller band seen in the FCCP-treated AIFM1^WT^ fraction contained the cleaved AIFM1 (Fig. 4F, middle blots). Importantly, this antibody did not cross-react with the AIFM1^TCS/TEV^ variant, which contains an additional glycine residue at the N-terminus due to TEV processing (Fig. 4A). Together, these results demonstrate that AIFM1 ordinarily forms larger complexes in the IMS (Hevler *et al*, 2021; Wang *et al*, 2021; Salscheider *et al*, 2022) that are modulated upon OMA1-mediated cleavage and the release of AIFM1 from the MIM.

### AIFM1 ensures cell proliferation by controlling OXPHOS activity

To define the AIFM1 assembly, we performed IP-MS to identify its interactome in mitochondria. Mitochondrial extracts from AIFM1^TCS^ and AIFM1^TCS/TEV^-expressing cells were subjected to IP, and the digested peptides were analyzed by MS (Fig. 5A and Dataset EV3). The proteomic data revealed that most of the enriched proteins in the AIFM1^TCS^ fraction belonged to two categories (Fig. 5A-B): subunits of the OXPHOS machinery and members of the mitochondrial carrier family (SLC25), both of which facilitate the transport of metabolites (Ruprecht & Kunji, 2020). Comparative analysis between these two variants revealed that components of mitochondrial complex I were enriched in the AIFM1^TCS^ fraction (Fig. 5C-D, highlighted in red [D]), supporting the finding that AIFM1 is required for complex I biogenesis (Vahsen *et al*, 2004; Urbano *et al*, 2005; Delavallée *et al*, 2020; Wang *et al*, 2021; Salscheider *et al*, 2022). We verified this result by Western blot analysis and confirmed that the AIFM1^TCS^, but not cleaved AIFM1^TCS/TEV^, was efficiently co-precipitated with some complex I, III, and IV components (Fig. 5E). These results clearly indicate that the assembly of AIFM1 with such interactors (a member of the complexome) is dependent on the MIM-anchored conformation.

**Figure 5.**
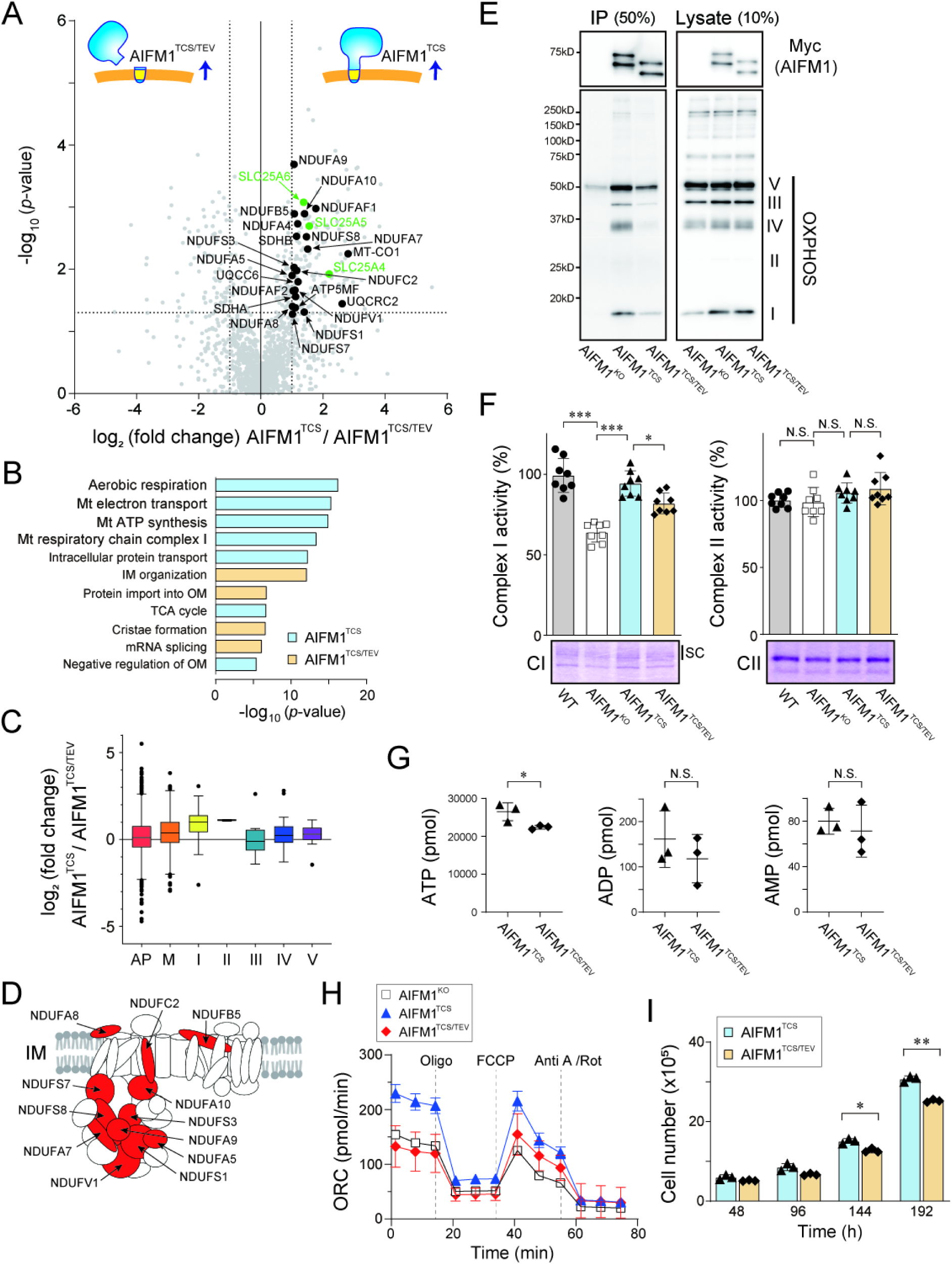
Functional interaction between AIFM1 and OXPHOS machinery. **A** Proteomic analysis to evaluate the interactome of AIFM1 dependent on membrane anchoring. Mitochondrial extracts from either formalin-fixed AIFM1^TCS^ or AIFM1^TCS/TEV^-expressing cells were immunoprecipitated. Co-purifying proteins were identified by quantitative MS (*n* = 3 biologic replicates) followed by statistical testing. Several subunits of OXPHOS (black) and members of the SLC25 family (green) enriched in the AIFM1^TCS^ fraction are indicated. See also Dataset EV3. **B** Ingenuity pathway analysis of IP-MS results from (A) showing the 11 most significantly altered pathways in AIFM1^TCS^ (blue) and AIFM1^TCS/TEV^ (orange). **C** Box and whisker plot showing log_2_ fold-change distributions of proteins quantified by LC-MS/MS in (A) that were identified in specific OXPHOS complexes I-V. AP = all detected proteins (1,397 proteins), M = MitoCarta3.0 (418 proteins), CI = complex I (39 proteins), CII = complex II (3 proteins), CIII = complex III (13 proteins), CIV = complex IV (34 proteins), and CV = complex V (12 proteins). The box borders indicate the 25% and 75% quantiles, and outliers are indicated by a black plot (greater distance than 1.5 times the interquartile range). **D** Illustration of the mitochondrial respiratory complex I, highlighting the positions of the 12 subunit positions with a high affinity for AIFM1, which are shown in red. **E** Interaction of OXPHOS components with AIFM1 variant. The *AIFM1* KO or AIFM1 variants (AIFM1^TCS^ and AIFM1^TCS/TEV^) expressing cells were moderately fixed with 0.2% formaldehyde and their lysates were immunoprecipitated with anti-Myc antibody followed by Western blotting analysis with an OXPHOS monoclonal antibody cocktail. Lysate and IP were loaded at 10% and 50% of the input samples, respectively. Complex I, NDUFB8; Complex II, SDHB; Complex III, UQCRC2; Complex IV, MTCO1; Complex V, ATP5A. **F** In-gel catalytic activity assay of digitonin-solubilized mitochondrial complex I and II in CN-PAGE (bottom image). In this assay, mitochondria were isolated from Flp-In-293 cells (WT, AIFM1 KO, AIFM1^TCS^, and AIFM1^TCS/TEV^). SC, Supercomplexes. Top, Each value represents the mean ± SD (*n* = 8 biologic replicates) and statistical analysis was performed by one-way ANOVA followed by Tukey’s test. \*\*\**p* < 0.001 and \**p* < 0.05. N.S., not significant. **G** ATP, ADP, and AMP levels (pmol/10^6^ cells) in AIFM1^TCS^ and AIFM1^TCS/TEV^ cells were analyzed by LC-MS/MS. Data shown are mean ± SD (*n* = 3 biologic replicates). \**p* < 0.05. N.S., not significant. **H** Oxygen consumption rate (OCR) of *AIFM1* KO or AIFM1 variants (AIFM1^TCS^ and AIFM1^TCS/TEV^) expressing cells. Graphs are mean ± SD of nine independent biologic experiments. Dashed lines indicate injections of oligomycin (2 μM), FCCP (0.5 μM), and antimycin A/rotenone (0.5 μM each). **I** Growth rate of the AIFM1 variant cells in a galactose-containing medium. Each cell was seeded at 4 × 10^5^ cells per well (in a 6-well plate) in a customized galactose-containing medium to switch the cellular biogenesis to rely on mitochondrial respiratory activity, and the number of cells was counted for up to 8 days after seeding. Data shown are mean ± SD (*n* = 3 biologic replicates). \*\**p* < 0.01 and \**p* < 0.05.

How does mislocalization of AIFM1 in mitochondria affect physiologic functions? The observed association of AIFM1 with OXPHOS components led us to investigate whether mislocalization of AIFM1 in mitochondria could affect its functional relevance in energy production via OXPHOS activity. To this end, we assessed the levels of complex I activity (i.e., NADH dehydrogenase activity) in cells using CN-PAGE (Wittig *et al*, 2007). Using this approach, *AIFM1* KO cells showed a significant reduction in complex I activity compared to WT cells (Fig. 5F), as previously reported (Delavallée *et al*, 2020; Salscheider *et al*, 2022). The attenuation of complex I activity observed in the *AIFM1* KO cells was substantially restored when the AIFM1^TCS^ variant was expressed in cells, but the restoration was not sufficient in the AIFM1^TCS/TEV^ cells (Fig. 5F). Consistent with these results, quantification of ATP levels in the AIFM1^TCS^ cells showed a much higher ATP concentration than in the AIFM1^TCS/TEV^ variant (Fig. 5G), and measurement of oxygen consumption rates also supported respiratory recovery in AIFM1^TCS^ cells (Fig. 5H and EV6D). These results indicate that AIFM1 anchoring to the MIM is correlated with control of the OXPHOS system. Most importantly, cell growth in galactose-containing medium, where cellular biogenesis would rely on mitochondrial respiratory activity (Mishra *et al*, 2014), was impaired in the AIFM1^TCS/TEV^ cells compared to the AIFM1^TCS^ cells (Fig. 5I). Taken together, these data demonstrate that the membrane-associated AIFM1 interactome ensures cell proliferation by controlling OXPHOS activity. Based on these results, we propose that OMA1 cleavage-mediated AIFM1 inactivation would lead to an OXPHOS deficiency.

### AIFM1 coordinately regulates substrate import by associating with TIM23 translocase

To substantiate these findings, we performed an MS-based analysis of the mitochondrial proteome in AIFM1-variant cells at steady state. The topologic change in AIFM1 was accompanied by extensive changes in the mitochondrial proteome (Fig. 6A and Dataset EV4). Gene Ontology annotation of the mitochondrial proteome in AIFM1^TCS/TEV^ cells (boxed region in Fig. 6A) revealed that mitochondrial respiration and the protein import pathway were predominantly affected (Fig. 6B), consistent with previous interactome results (Fig. 5). Therefore, we compared the enrichment profile of mitochondrial proteins found in the AIFM1^TCS/TEV^ variant cells. Although membrane dislocation of AIFM1 did not grossly alter mitochondrial DNA levels (Fig. EV6E), expression of OXPHOS-related genes (Fig. EV6F), or the ΔΨ_m_ (Fig. EV6G), 32 subunits of the OXPHOS complex were decreased in the AIFM1^TCS/TEV^ variant (Fig. EV6H). These results imply that membrane-associated AIFM1 maintains the protein levels of each OXPHOS subunit at steady state.

**Figure 6.**
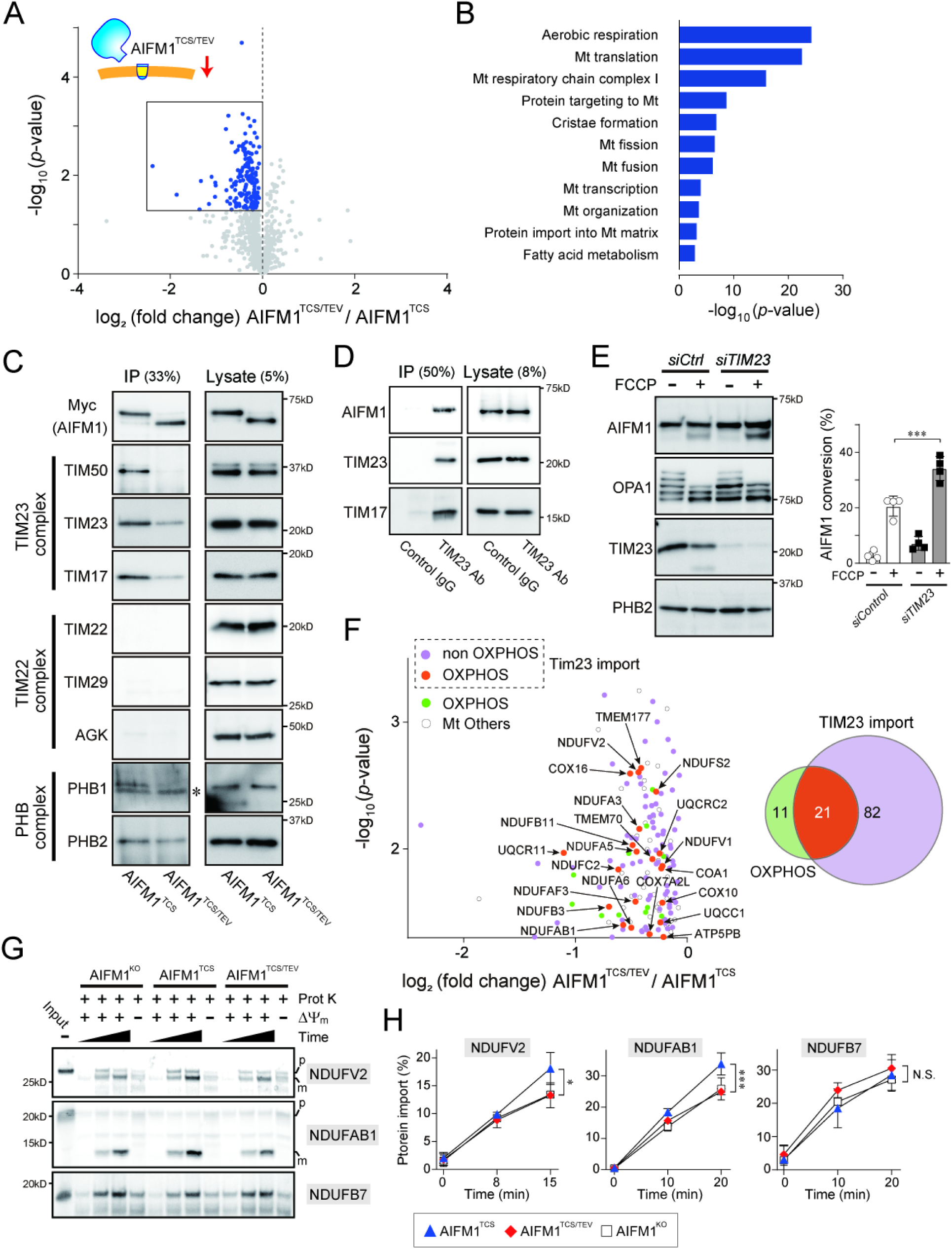
AIFM1 cooperates with TIM23 translocase and ensures substrate import. **A** Volcano plot of mitochondrial protein changes (MitoCarta3.0) between AIFM1^TCS^ and AIFM1^TCS/TEV^ cells. Blue plot area in the volcano plot shows decreased proteins (*p* < 0.05, *n* = 4 biologic replicates) in AIFM1^TCS/TEV^, and several mitochondrial proteins in the boxed area are annotated in (F). See also Dataset EV4. **B** Ingenuity pathway analysis of the mitochondrial proteome from the boxed area in (A), showing significantly altered pathways in AIFM1^TCS/TEV^. **C** Mitochondrial extracts from the AIFM1 variant cells were immunoprecipitated with anti-Myc antibody and subjected to immunoblotting using the indicated antibodies. Lysate and IP samples were loaded at 5% and 33% of the input samples, respectively. **D** Interaction of endogenous AIFM1 with TIM23 translocase. Lysates of HeLa cells were immunoprecipitated with either anti-TIM23 monoclonal antibody or control IgG followed by Western blot analysis with the antibody against AIFM1. The lysate and IP samples were loaded at 8% and 50% of the input samples, respectively. As for a control, we confirmed that endogenous TIM23 was immunoprecipitated with the component of the TIM23 translocase, TIM17. **E** HeLa cells transfected with siRNA against *TIM23* were treated for 3 h with or without FCCP (40 μM) and analyzed by immunoblotting as indicated. The graph on the right shows the quantification of AIFM1 bands from the immunoblot analyzed by densitometry. **F** Enlarged box area in (A). A total of 123 mitochondrial proteins are plotted, 103 of which are TIM23 substrates, including the 21 OXPHOS components (orange). See also Dataset EV4. **G** *In organello* import assay with TIM23 substrates (NDUFV2 and NDUFAB1). *In vitro* radiolabeled NDUFV2 or NDUFAB1 (^35^S-labeled) were incubated with mitochondria isolated from the *AIFM1* KO cells or cells expressing AIFM1 variant (AIFM1^TCS^ and AIFM1^TCS/TEV^). Non-imported proteins were removed by treatment with proteinase K (Prot K), and imported proteins at different time points were analyzed by SDS-PAGE followed by autoradiography (top). As a control, the import reaction was also performed on mitochondria treated with uncouplers to dissipate the ΔΨ_m_. Radioactive NDUFB7 was used as a control for TIM23-independent substrate in this assay. NDUFV2 and NDUFAB1 import is affected by the loss or membrane dislocation of AIFM1, but NDUFB7 import is less affected. *p*, precursor form; *m*, mature form. **H** Signals in (G) were quantified using ImageJ and the amount of each imported protein at different time points was plotted. Data shown are mean ± SD (*n* = 3 biologic replicates). \**p* < 0.05, \*\*\**p* < 0.001, and N.S, not significant, respectively. Each statistic compares AIFM1^TCS^ vs AIFM1^TCS/TEV^ and AIFM1^TCS^ vs. AIFM1^KO^, respectively.

To explore the possible involvement of AIFM1 in the biogenesis of OXPHOS subunits, we next examined whether AIFM1 could maintain its protein import by cooperating with import machinery (Salscheider *et al*, 2022; Peker *et al*, 2023). To this end, we investigated the physical interaction between AIFM1 and mitochondrial translocase complexes by IP assay. Using the AIFM1 variant cells, we found that AIFM1^TCS^ interacted with subunits consisting of the translocase of the inner membrane 23 (TIM23) complex (TIM23, TIM50, and TIM17), but not with the TIM22 translocase (Fig. 6C). By contrast, the ability of AIFM1^TCS/TEV^ to bind to the TIM23 translocase was weakened, as also confirmed by the IP-MS result (Fig. EV6I). The membrane-associated AIFM1 variant also showed substance affinity for the MIM scaffold, the PHB complex (particularly for PHB1) (Fig. 6C). Consistent with these results, we confirmed that TIM23 also co-immunoprecipitated endogenous AIFM1 in HeLa cells (Fig. 6D). Intriguingly, the physical interaction between the AIFM1 and TIM23 translocase affected the susceptibility of AIFM1 to OMA1-dependent proteolytic processing. When TIM23 was depleted from cells by siRNA, OMA1-mediated AIFM1 processing was significantly increased under depolarized mitochondria (Fig. 6E), suggesting that OMA1 accessibility to AIFM1 was increased by removal of the TIM23 complex.

Having identified that the membrane-targeted AIFM1 is associated with the TIM23 translocase in mitochondria, we revisited the enrichment analysis of the mitochondrial proteome in AIFM1 variant cells and sought to identify related molecules that rely on the TIM23 import pathway. LC-MS/MS identified 114 individual proteins that were downregulated in the AIFM1^TCS/TEV^ cells (Fig. 6A, boxed region), and defined the majority of these (103 proteins) as substrates of the TIM23 translocase (Crameri *et al*, 2024) (Dataset EV4). In particular, we found that 21 of the 103 mitochondrial proteins correspond to the subunits of the OXPHOS complex that are imported via the TIM23 pathway (Fig. 6F, orange). Indeed, our *in vitro* import assay revealed that loss of AIFM1 or its membrane anchoring resulted in moderately impaired protein import of NDUFV2 or NDUFAB1, the 24-kDa or 16-kDa subunits of complex I, into mitochondria compared with AIFM1^TCS^-containing mitochondria, but not the TIM23-independent substrate NDUFB7 (Crameri *et al*, 2024) (Fig. 6G-H). We conclude that AIFM1 anchoring to the MIM results in a functional interaction with the TIM23 translocase, which coordinately regulates protein import into mitochondria.

## Discussion

OMA1 is a stress-sensitive mitochondrial metalloprotease that belongs to the membrane-integrated peptidase M48 superfamily (López-Pelegrín *et al*, 2013). It plays diverse roles in cellular processes throughout the organism, although its catalytic mechanisms, especially how it recognizes and processes substrates, remain poorly understood. In the present study, we used multiproteomic and biochemical approaches to determine the interactome of the OMA1 peptidase in mitochondria and identified AIFM1 as an OMA1-targeted substrate under various mitochondrial stresses. OMA1-mediated cleavage of AIFM1 leads to membrane dislocation of the substrate and downregulates the import of OXPHOS-related proteins into mitochondria, ultimately resulting in impaired mitochondrial bioenergetics.

OMA1-mediated AIFM1 processing is strictly triggered by mitochondrial stress. Notably, OMA1 substrates do not share a conserved cleavage motif (Fig. EV3B) (Ishihara *et al*, 2006; Guo *et al*, 2020; Ahola *et al*, 2024), and the cleavage site in AIFM1 does not match those targeted by calpain or cathepsins during apoptosis (Susin *et al*, 1999; Polster *et al*, 2005; Yuste *et al*, 2005). Our biochemical data show that the OMA1-mediated AIFM1 processing kinetics are significantly slower than those of the conventional substrate, OPA1 (Fig. 3B). Like AIFM1 proteolysis, the kinetic profile of OMA1-induced DELE1 cleavage is also slower (Fessler *et al*, 2020), although there is no apparent interaction between these two substrates. Because elimination of OPA1, but not DELE1 (Fig. EV5D), in cells would accelerate OMA1-mediated processing of AIFM1 (Fig 3E), we believe that the OMA1 peptidase has substrate selectivity and that OPA1 has a much higher affinity than other substrates, which is also supported by the fact that OPA1 is constitutively processed at steady state (Ishihara *et al*, 2006; Song *et al*, 2007; Ehses *et al*, 2009; Head *et al*, 2009). This may explain why AIFM1 cleavage is observed in depolarized *OPA1-*depleted cells at early time points, increasing its susceptibility to OMA1 attack (Fig. EV5B). Although the catalytic reaction of OMA1 in AIFM1 processing appears to act in *cis* (i.e., two molecules located on the same membrane), we cannot exclude the possibility that each molecule exists on a different membrane (*trans*) and proteolysis occurs when the membranes come closer together (Fig. EV2E), allowing the molecules to persist for a longer period of time. Together, this unusual catalytic mechanism of OMA1 regulates a rapid change in mitochondrial dynamics when stress stimuli are induced, while subsequently leading to an alternative stress response program in cells, ensuring functional plasticity at the cellular level.

We found that AIFM1 functionally associates with the TIM23 complex, which coordinately regulates the translocase-dependent import of substrates into the matrix. The function of TIM23 is critically dependent on ATP hydrolysis and the ΔΨ_m_ (Neupert & Brunner, 2002; Chacinska *et al*, 2009; Mokranjac & Neupert, 2010; van der Laan *et al*, 2010), so direct investigations of the mechanistic action of AIFM1 involved in the TIM23-dependent import pathway under the ΔΨ_m_-less condition have been challenging. Our platform, which generated the AIFM1 cleaved protein without inducing any mitochondrial stress (e.g., depolarization) overcomes these limitations and allowed us to reveal the regulatory role of AIFM1 involved in TIM23-mediated mitochondrial protein import. Our results suggest that release of AIFM1 from MIM attenuates some interactions between AIFM1 and the TIM23 complex, and that this dissociation ultimately downregulates the mitochondrial proteome and impairs cell growth. Indeed, the membrane dislocation of AIFM1 decreased protein levels of 32 OXPHOS-related components at steady state, with >60% of them being TIM23-dependent pathways (Fig. 6F). These results are consistent with previous observations implicating AIFM1 in OXPHOS biogenesis (Vahsen *et al*, 2004; Urbano *et al*, 2005; Delavallée *et al*, 2020; Wang *et al*, 2021; Salscheider *et al*, 2022; Peker *et al*, 2023) and support the possibility that AIFM1 functions as a chaperone activity involved in regulating OXPHOS metabolism (Brosey *et al*, 2025). Accordingly, these results provide a view of AIFM1 as an essential import platform not only for components of the mitochondrial disulfide relay (Reinhardt *et al*, 2020; Salscheider *et al*, 2022; Peker *et al*, 2023; Brosey *et al*, 2025), but also for OXPHOS components translocated into the matrix. AIFM1 mutations are reported in humans with autosomal recessive mitochondrial disorders (Bano & Prehn, 2018; Wischhof *et al*, 2022), although their precise physiologic functions with biochemical properties of the mutations remain elusive. While these *AIFM1* mutation-associated disorders manifest variable clinical manifestations, many of the *AIFM1* mutant alleles generally exhibit respiratory defects (Wischhof *et al*, 2022). It is interesting to consider our results with previous findings that two AIFM1 variants (deletion of residue Arg^201^; R201 del, and substitution of residue Glu 493 to Val; E493V), both of which fold nearly identically to the WT protein, exhibit higher sensitivity to tryptic digestion *in vitro* (Ghezzi *et al*, 2010; Rinaldi *et al*, 2012), suggesting that subtle conformational changes due to the removal or substitution of residues in AIFM1 destabilize the stability of the protein, making it more susceptible to proteolytic attack. Indeed, Western blot analysis of three alleles of patient fibroblasts reveals an apparent reduction in AIFM1 abundance (Ardissone *et al*, 2015; Diodato *et al*, 2016; Moss *et al*, 2021). It remains to be directly demonstrated that disease-associated mutations (or deletion) in AIFM1 increase protein destabilization, but the present observations raise the possibility that some mitochondrial proteases are involved in the degradation process, leading to AIFM1 inactivation in mitochondrial bioenergetics, and that OMA1 may act to fulfill this role. Therefore, additional investigation into how OMA1 functionally interacts with other mitochondrial proteases to participate in the degradation process of AIFM1 variants and their impact on human disease will provide further elucidation.

## Materials and Methods

### Reagents and tools

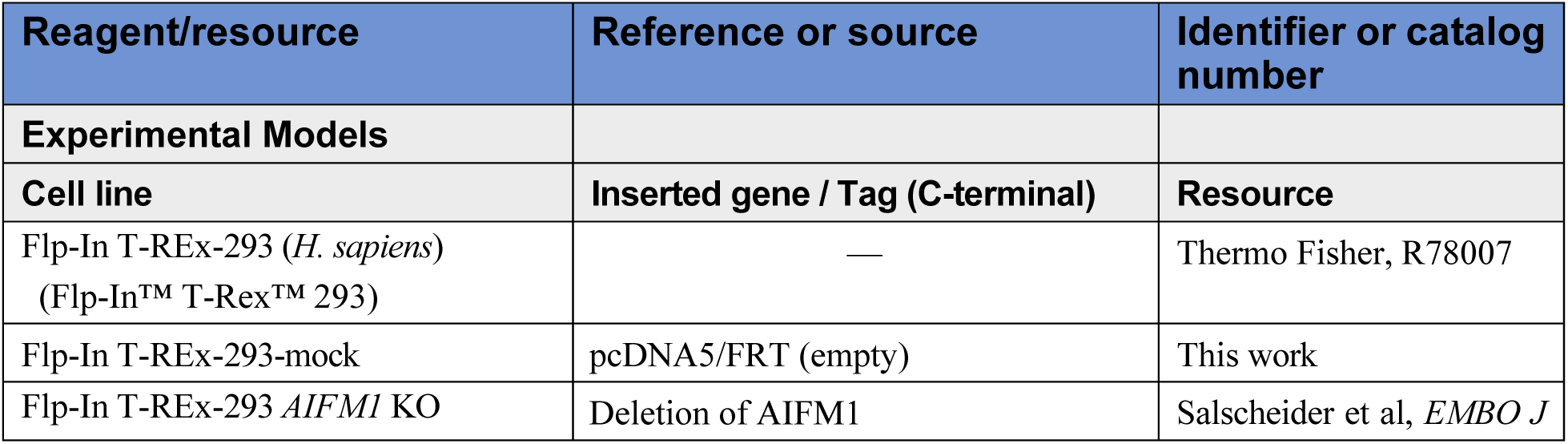
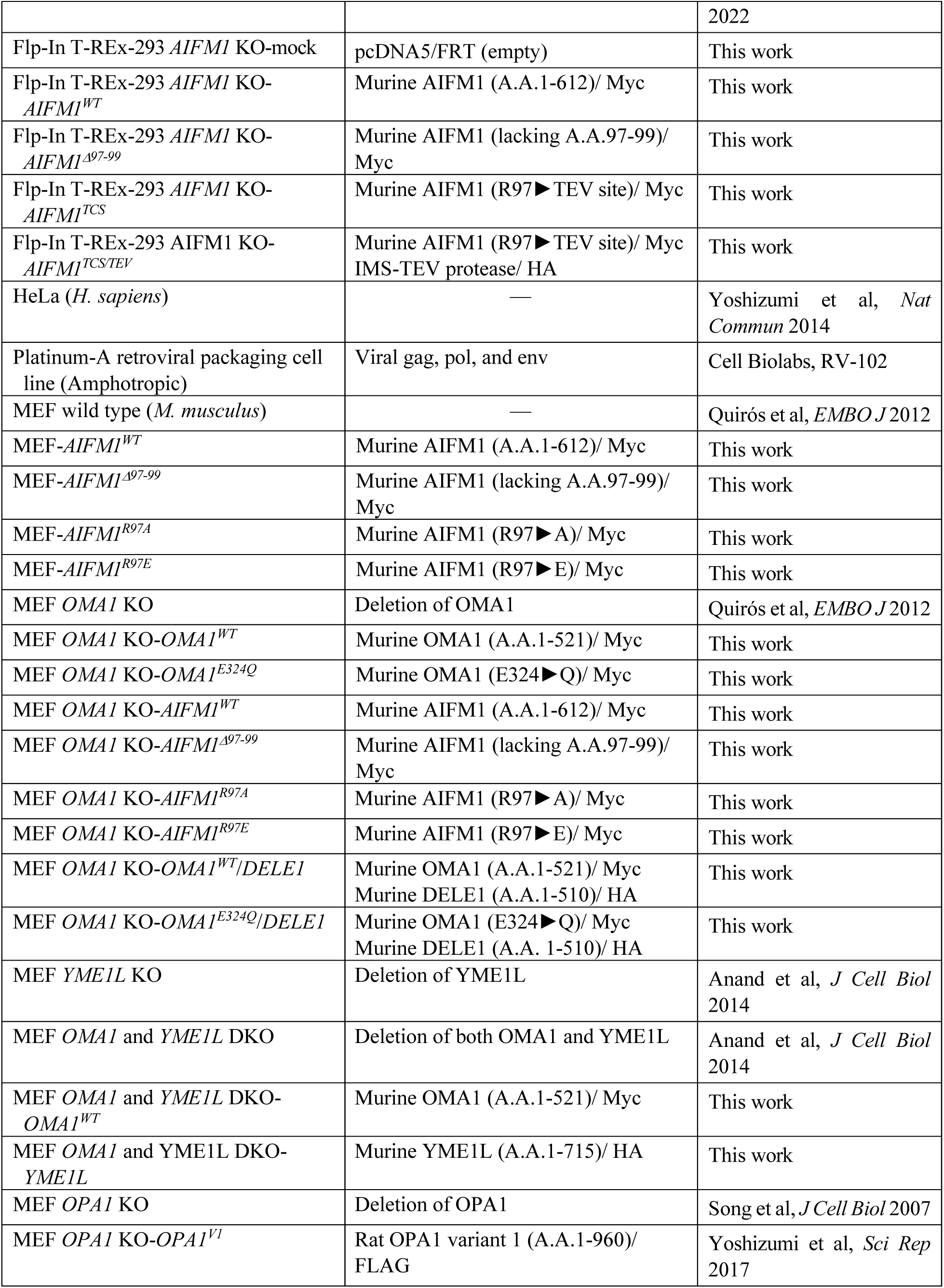
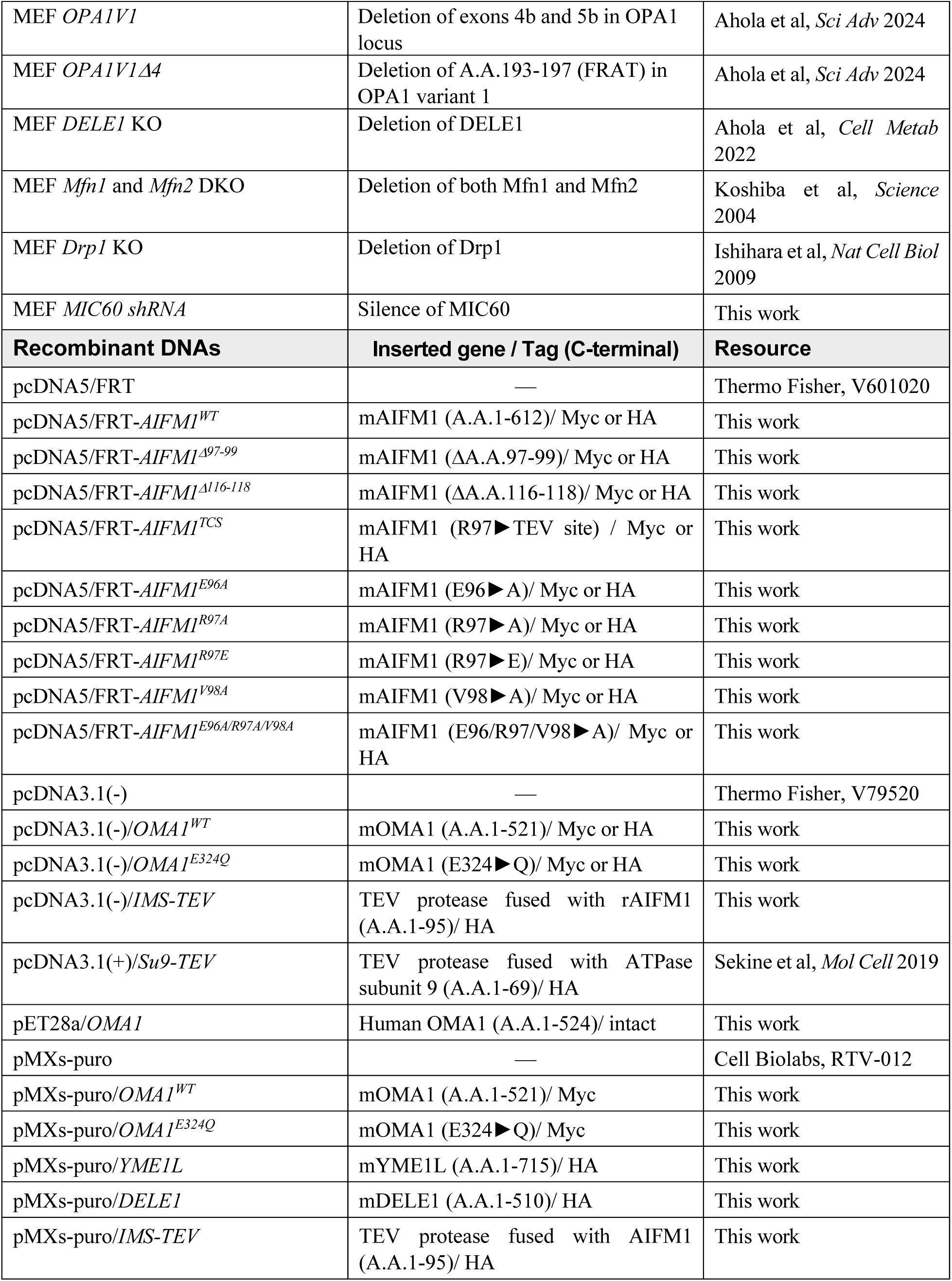
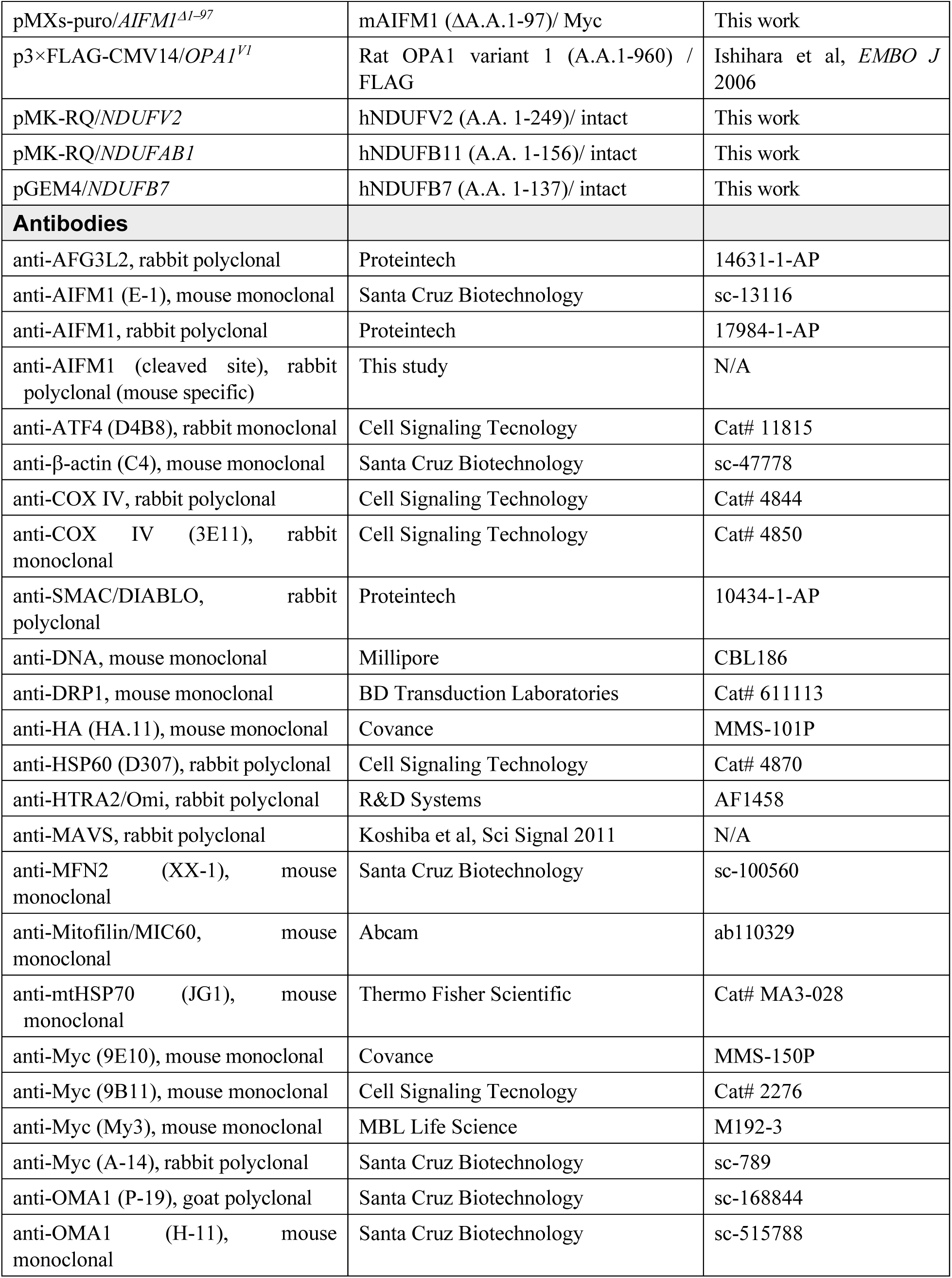
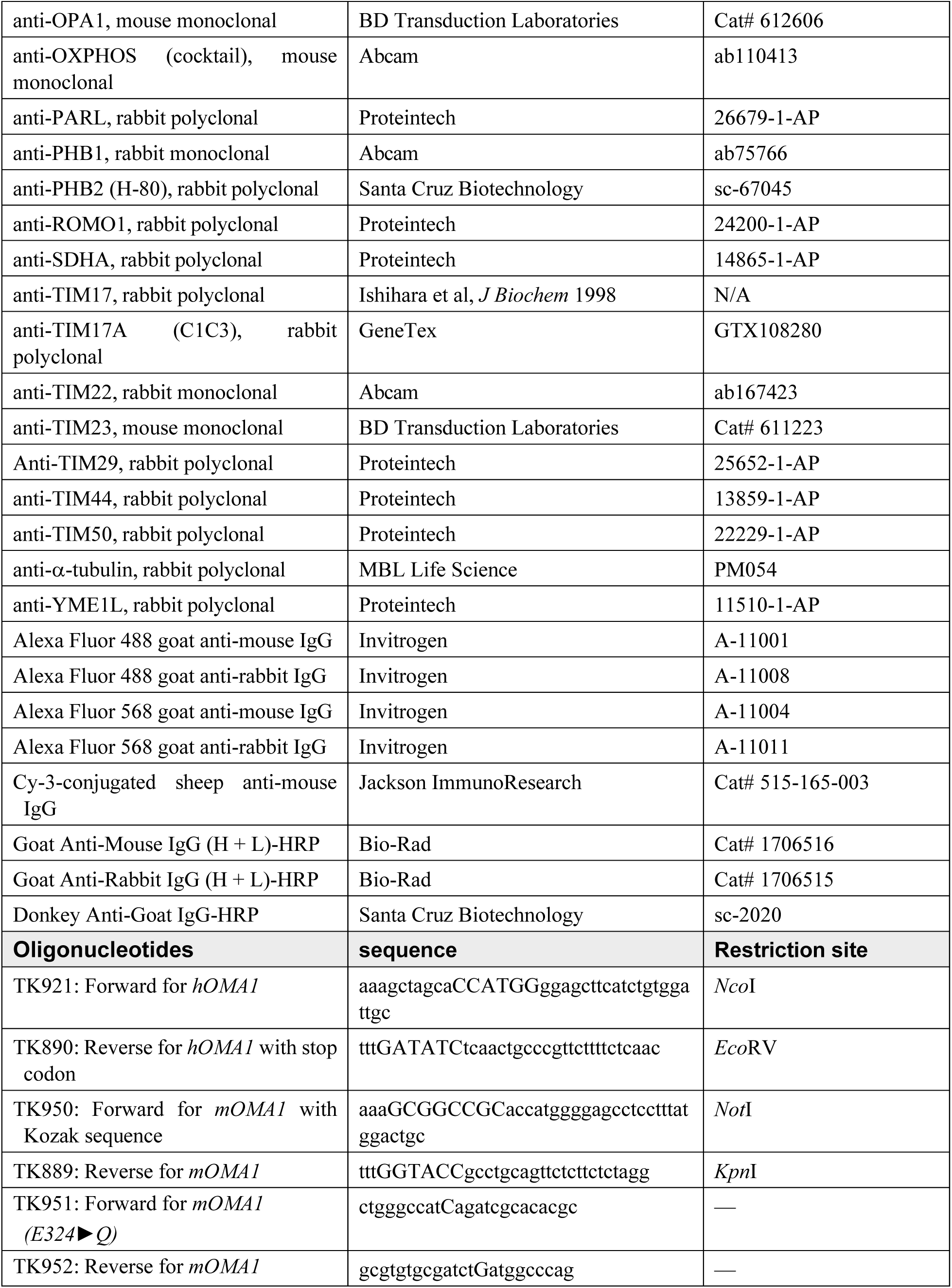
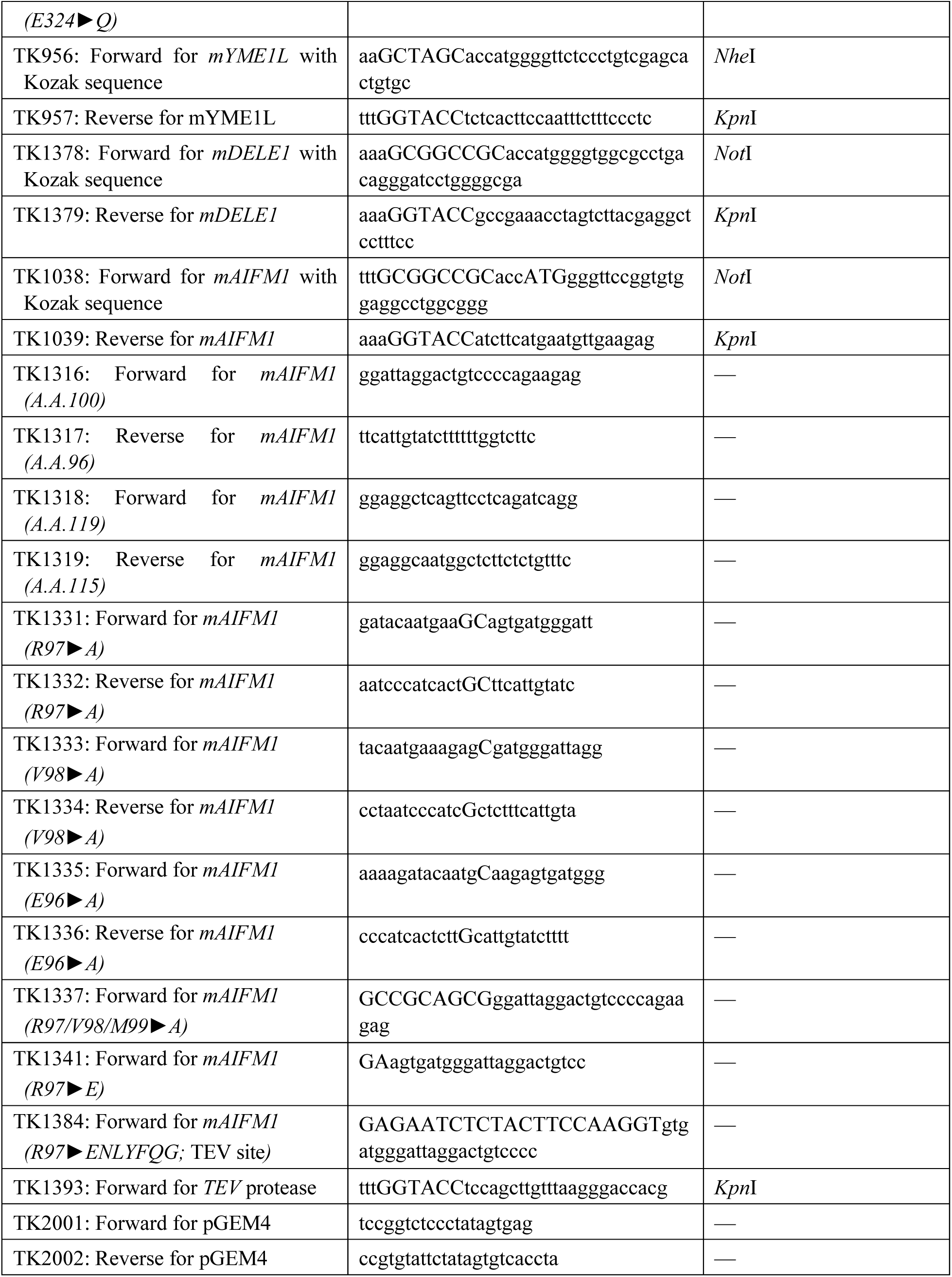
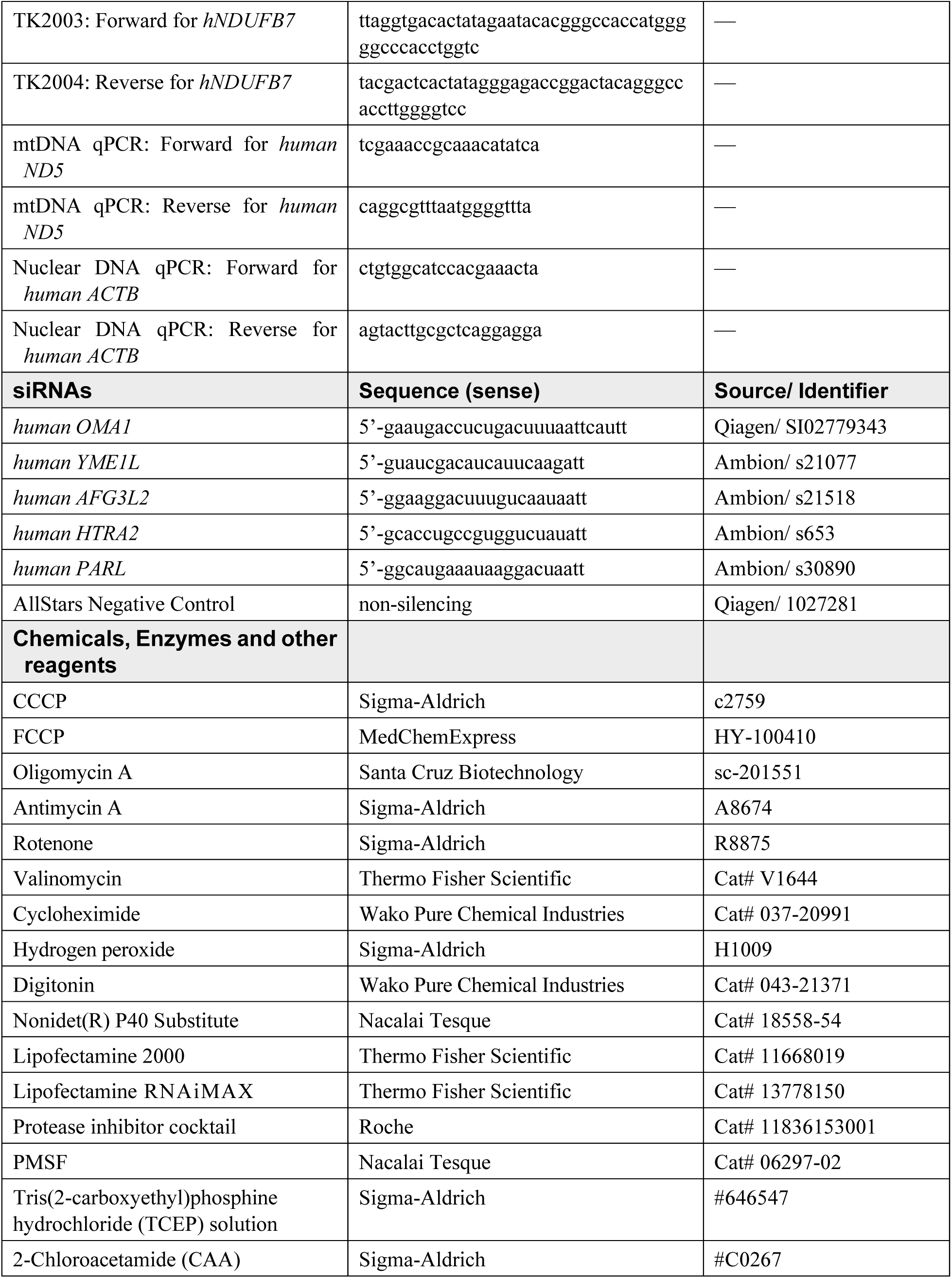
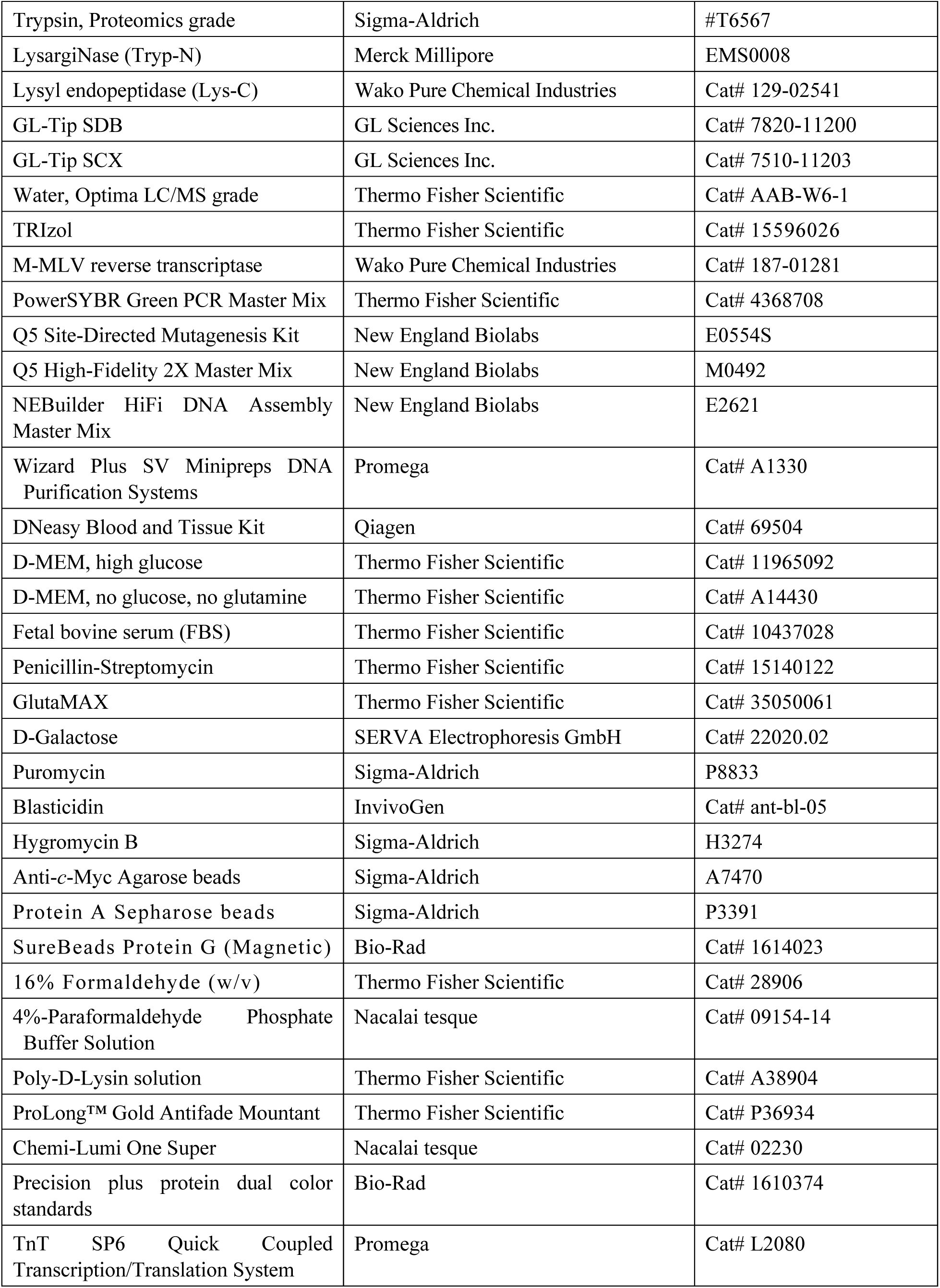
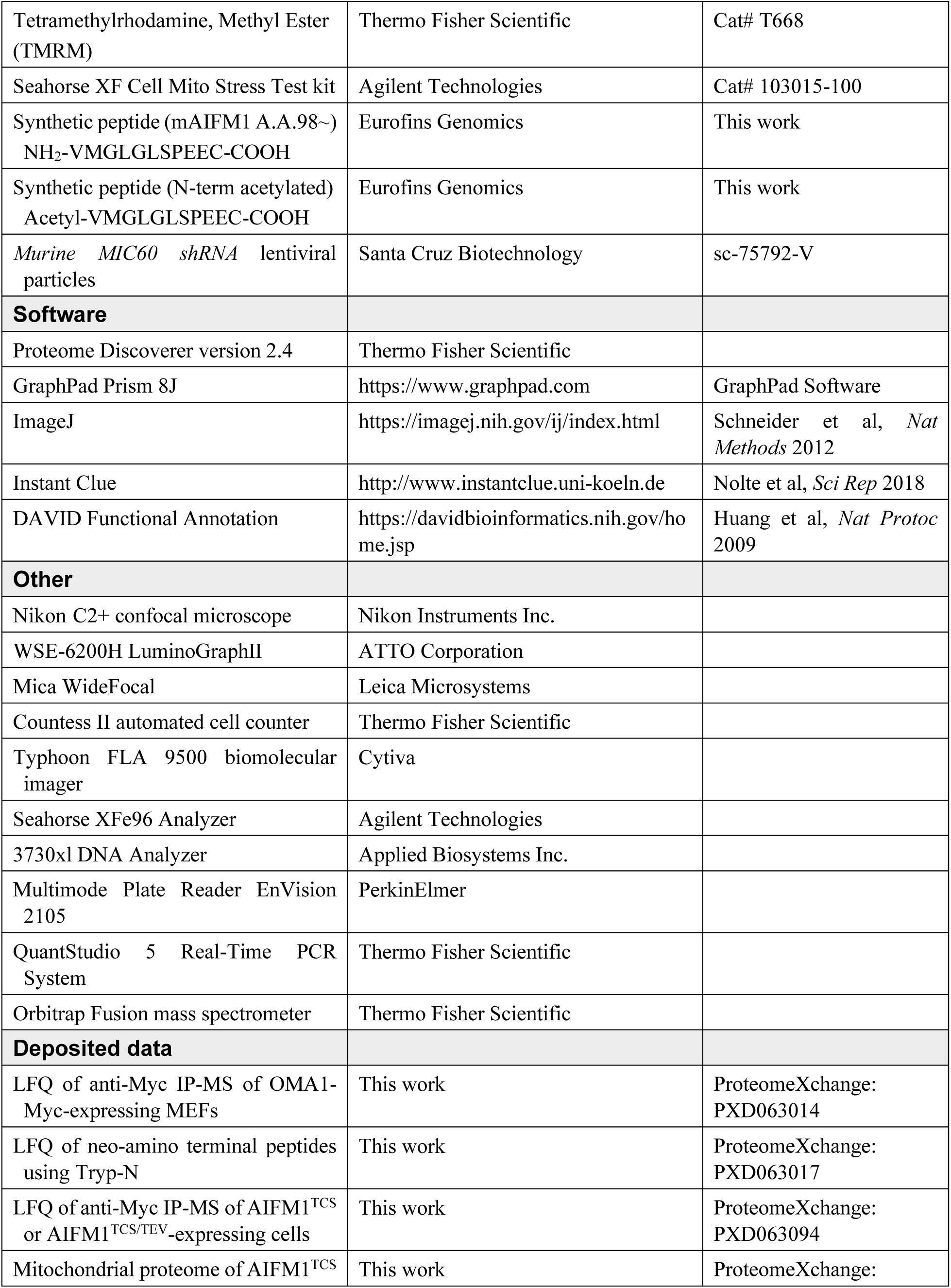
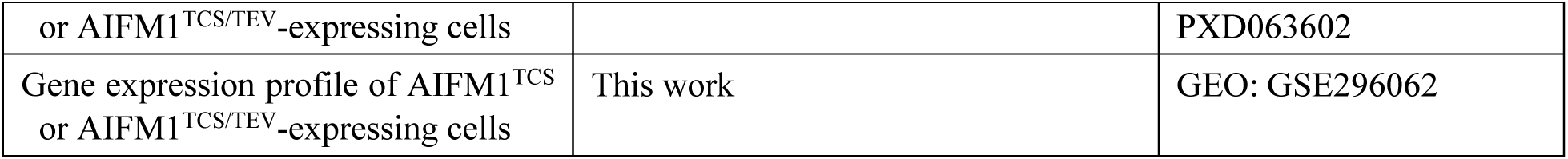

#### Methods and protocols Reagents

CCCP (Cat# c2759), antimycin A (Cat# A8674), rotenone (Cat# R8875), hygromycin B (Cat# H3274), puromycin (Cat# P8833), hydrogen peroxide (Cat# H1009), anti-*c*-Myc agarose beads (Cat# A7470), and Protein A Sepharose beads (Cat# P3391) were purchased from Sigma Aldrich (St. Louis, MO). FCCP (Cat# HY-100410) and oligomycin A (sc-201551) were supplied from either MedChemExpress (Monmouth Junction, NJ) or Santa Cruz Biotechnology (Dallas, TX). Valinomycin (Cat# V1644) and cycloheximide (Cat# 037-20991) were obtained from Thermo Fisher Scientific (Waltham, MA) and Wako Pure Chemical Industries (Tokyo, Japan), respectively. All other reagents used in the study were biochemical research grade.

#### Cell culture and cell treatments

All cell lines used in the study are listed in the Reagents and tools Table. HeLa cells were used for siRNA-mediated knockdown experiments and the Platinum-A retroviral packaging cell line (Cell Biolabs, San Diego, CA) was used for retrovirus production. Unless otherwise indicated, cells were maintained in Dulbecco’s modified Eagle medium (D-MEM, high glucose) supplemented with 1% GlutaMAX, penicillin (100 U/mL)-streptomycin (100 μg/mL), and 10% fetal bovine serum (FBS, Thermo Fisher Scientific) at 37° C under 5% CO_2_ conditions. Mitochondrial stress was induced by treating cells with CCCP (20 μM), FCCP (20 μM), valinomycin (1 μg/mL), oligomycin A (1 μM), antimycin A (1 μM), H_2_O_2_ (1 mM), or rotenone (5 μM) as indicated. In some experiments, cycloheximide (10 μg/mL) was added to the culture medium to inhibit protein synthesis. Cells were regularly seeded at equal densities before an experiment and checked for mycoplasma contamination.

##### Plasmid construction and mutagenesis

Total mRNAs from HEK293 and MEFs were isolated using TRIzol reagent (Thermo Fisher Scientific) and reverse transcribed using M-MLV reverse transcriptase (Wako Pure Chemical Industries). Polymerase chain reaction assays were performed using Q5 High-Fidelity DNA polymerase (New England Biolabs, Ipswich, MA). The following primers (see Reagents and tools Table for sequences) were used to generate the complete open reading frames of human OMA1: TK921/TK890; murine OMA1: TK950/TK889; mYME1L: TK956/TK957; mDELE1: TK1378/ TK1379; mAIFM1: TK1038/TK1039. Plasmids encoding epitope-tagged proteins were constructed by ligation of each cDNA into either *Not*I or *Nhe*I at the 5’ end and *Kpn*I at the 3’ end of a digested pcDNA5/FRT vector (Thermo Fisher Scientific) encoding either a C-terminal 3× Myc or 3× hemagglutinin (HA) tag. To generate a plasmid encoding TEV protease localized to the mitochondrial IMS (IMS-TEV), a DNA fragment encoding the IMS targeting sequence of rat AIFM1 (A.A.1-95) was cloned in front of the sequence encoding the TEV protease. The substitution was introduced into each plasmid by site-directed mutagenesis according to the manufacturer’s protocol (New England Biolabs). To generate retroviral expression constructs, each cDNA was subcloned into the retroviral vector pMXs-puro (Cell Biolabs). All constructs used in the study were confirmed by DNA sequencing (Applied Biosystems 3730xl Genetic Analyzer, Waltham, MA).

##### Generation of stable expression cell lines

Flp-In T-REx-293 AIFM1 knockout HEK293 cells (Salscheider *et al*, 2022) were transfected with either pcDNA5/FRT-AIFM1^WT^ or AIFM1 variants. After transfection, cells were selected with 100 μg/mL hygromycin B for 7-10 days, single colonies were selected by plating, and expression of the gene of interest at the steady state levels was checked by Western blotting. Viral gene delivery using a retrovirus system was also used to generate stable expression cell lines. Briefly, each constructed pMXs-puro plasmid (see Reagents and tools Table) was transfected into the platinum packaging cell lines using Lipofectamine 2000 reagent (Thermo Fisher Scientific), and then the retroviral supernatants were harvested 48 h post-transfection and used to infect MEF or other cell lines. After retroviral infection, cells were selected with 1.5 μg/mL puromycin for 1 week, and single colonies were selected by checking expression by immunoblotting. All antibodies used in the study are listed in the Reagents and tools Table.

#### Protein expression in *Escherichia coli*

Recombinant human OMA1 was expressed in *E. coli* BL21(DE3) cells (Novagen, Madison, WI). The inoculated culture was grown to log phase at 37° C, and overproduction of the recombinant protein was induced by the addition of isopropyl β-D-1-thiogalactopyranoside (Nacalai Tesque, Kyoto, Japan) to a final concentration of 1 mM. After 3 h of induction, cells were harvested by centrifugation (6,000*g* for 15 min), and the pellets were stored frozen (−20° C) until used for the Western blotting.

#### Mitochondrial isolation and proteolysis

Mitochondrial isolation, proteolysis, and membrane association assays were performed as previously described (Yoshizumi *et al*, 2014) with slight modifications. Briefly, cultured cells were washed once with 1× phosphate-buffered saline (PBS, pH 7.4), scraped from the culture plate, and lysed in ice-cold homogenization buffer containing 20 mM HEPES (pH 7.5), 70 mM sucrose, and 220 mM mannitol by 30 strokes in a Dounce homogenizer on ice. The homogenate was then centrifuged at 800*g* for 5 min (4° C) to precipitate the nuclei, and the resulting supernatant was further centrifuged at 10,000*g* for 10 min (4° C) to precipitate the crude mitochondrial fraction.

For the proteinase K resistance assay, the isolated mitochondrial pellet was resuspended in either the homogenization buffer or a hypotonic buffer (20 mM HEPES, pH 7.5) and kept on ice for 30 min to cause swelling. Both samples were then treated with proteinase K (50 μg/mL) for 15 min on ice, and the reactants were subjected to Western blot analysis with the indicated antibodies. For the membrane association assay, mitochondrial pellets with or without swelling treatment were washed once with the homogenization buffer and centrifuged at 15,000*g* for 15 min (4° C) to separate supernatant or precipitate fractions. The supernatant was subjected to TCA precipitation, and the precipitated fraction was resuspended in a sodium dodecyl sulfate (SDS) sample buffer (50 mM Tris-HCl [pH 6.8], 2% SDS, 0.1 % [w/v] bromophenol blue, 10% [v/v] glycerol, and 5% β-mercaptoethanol) and analyzed by Western blotting.

#### Generation of the N-terminal specific antibody against AIFM1

To generate a rabbit polyclonal antibody against the N-terminal region of the murine AIFM1 cleavage site (A.A. ∼98), antiserum raised against the synthetic peptide NH_2_-^98^VMGLGLSPEEC-COOH (Eurofins Genomics, Louisville, KY) was affinity-purified using the same synthetic peptide column. The purified fraction was then passed through a column with the N-terminally acetylated peptide sequence (CH_3_CO-NH_2_-^98^VMGLGLSPEEC) to remove immunoglobulin G (IgG), which recognizes the peptides mimicking the uncleaved protein. Finally, the flow-through fraction was affinity-purified on Protein A Sepharose (Sigma Aldrich).

#### Western blotting

Immunoblotting was performed as previously described (Koshiba & Chan, 2003) with minor modifications. Briefly, cells were collected and washed with 1× PBS (pH 7.4) before lysis with ice-cold lysis buffer (50 mM Tris-HCl buffer [pH 7.4] containing 150 mM NaCl, 1 mM EDTA, 1% [w/v] NP-40, 10% [w/v] glycerol and protease inhibitor cocktail; Roche, Basel, Switzerland). The extracts were then mixed with the SDS sample buffer and samples were separated by SDS-PAGE. Precision plus protein dual color standards (Bio-Rad, Hercules, CA) were used to estimate the molecular weight of each protein of interest on SDS gels. Immunoblotting of the gels was performed on Immobilon-P polyvinylidene difluoride (PVDF) membrane (Merck Millipore, Burlington, MA) for 60 min at 25 V using transfer buffer (23 mM Tris, 180 mM glycine, 19% methanol). Immunoblotted membranes were then blocked in 5% nonfat dry milk in Tris-buffered saline (TBS) containing 0.1% Tween-20, followed by incubation with the indicated primary antibodies, and proteins were detected with horseradish peroxidase-conjugated secondary antibodies using a WSE-6200H LuminoGraphⅡ Image Analyzer (ATTO Corporation, Tokyo, Japan). All antibodies used in the study are listed in the Reagents and tools Table.

#### RNA interference

For RNA interference knockdown experiments, siRNAs listed in the Reagents and tools Table were used. HeLa cells were each transfected twice with 10 nM siRNA (final concentration) at 48 h intervals using Lipofectamine RNAiMAX reagent (Thermo Fisher Scientific) according to the manufacturer’s protocols. At 96 h after the first treatment, the siRNA-treated cells were used for some functional assays. The AllStars Negative Control siRNA (Qiagen, Hilden, Germany) was used as a control.

#### Immunofluorescence

Cells were plated on coverslips in 12-well plates (5 × 10^4^ cells/well). For monitoring the Flp-In T-REx-293 cell line, the coverslips were pre-coated with 0.05 mg/mL poly-D-lysin solution (Thermo Fisher Scientific) before seeding the cells. The next day, the cells were fixed with 4% paraformaldehyde phosphate buffer solution (Nacalai Tesque) for 10 min at 37° C, permeabilized with 0.2% Triton X-100 in 1× PBS (pH 7.4), and blocked with 5% FBS. Epitope-tagged proteins (HA- or Myc-) were detected with their specific primary and AlexaFluor488 secondary antibodies (Invitrogen, Waltham, MA), and mitochondria were stained with either anti-COX IV (3E11) or mtHSP70 (JG1) primary antibodies followed by the AlexaFluor568 or Cy3-conjugated secondary antibodies, respectively. Cells were imaged using a C2+ confocal (Nikon Instruments Inc., Melville, NY) or Mica WideFocal (Leica Microsystems, Wetzlar, Germany) microscope.

#### Immunoprecipitation

For the immunoprecipitation experiment, ∼95% confluence cells stably expressing Myc-tagged AIFM1 constructs were washed once with 1× PBS (pH 7.4), lysed with lysis buffer containing 50 mM Tris-HCl (pH 7.4), 150 mM NaCl, 10% (w/v) glycerol, 1 mM EDTA, and 0.5% (w/v) digitonin, protease inhibitor cocktail, and the clarified supernatants were incubated with 1 μg of the anti-*c*-Myc monoclonal antibody 9E10 (Covance), followed by incubation overnight at 4° C with 20 μL Protein A-Sepharose beads (Sigma-Aldrich). The next day, the beads were washed four times with 1× PBS (pH 7.4), the precipitated proteins were eluted from the beads by adding SDS sample buffer and heating to 96° C for 3 min, and the immunoprecipitates were resolved by SDS-PAGE and immunoblotted with the indicated antibodies. For immunoprecipitation of the OXPHOS complex, samples were heated to 50° C for 10 min to avoid protein degradation.

#### Clear native-PAGE analysis

CN-PAGE was performed as previously described (Ban *et al*, 2025) with slight modifications. Briefly, isolated mitochondria from each cell were solubilized in 20 mM Bis-Tris (pH 7.0) buffer containing 2 mM NaCl, 500 mM aminocaproic acid, 10% (w/v) glycerol, 1 mM EDTA, 0.5% (w/v) digitonin, and protease inhibitor cocktail for 15 min on ice. The clarified supernatant was then supplemented with 0.5% (w/v) CBB-G250 and separated on a 3%-12% gradient gel. Anode buffer (50 mM Bis-Tris, pH 7.0) and cathode buffer (50 mM Tricine, 15 mM Bis-Tris [pH 7.0], 0.05% sodium deoxycholate, and 0.02% *n*-dodecyl-D-maltoside) were used for electrophoresis. After electrophoresis, the separated proteins in the gel were denatured at 60° C for 15 min in Tris-HCl (pH 6.8) buffer containing 1% SDS and 2% β-mercaptoethanol, transferred to a PVDF membrane, followed by Western blotting analysis.

#### LC-MS/MS analysis (IP-MS & Neo-amino terminal MS)

##### IP-MS

Control parental *OMA1* KO MEFs and rescued cells stably expressing WT OMA1-Myc (cultured in 10-cm dishes) were incubated with 0.2% (w/v) formaldehyde (Thermo Fisher Scientific) for 15 min at 37° C, followed by quenching with 100 mM glycine-NaOH (pH 7.5) for 10 min at room temperature. After washing once with 1× PBS (pH 7.4), the cells were scraped, and mitochondria were isolated from the cells as described above. The mitochondrial fractions were then lysed in 1 mL of lysis buffer containing 50 mM Tris-HCl (pH 7.4), 150 mM NaCl, 10% (w/v) glycerol, 1 mM EDTA, 0.5% (w/v) digitonin, and protease inhibitor cocktail. After centrifugation, the clarified supernatants were incubated with an antibody against Myc (My3; MBL Life Science, Tokyo, Japan) for 2 h at 4° C. The reactants were then incubated with magnetic SureBeads Protein G (Bio-Rad) overnight at 4° C, and the next day the beads were washed three times with 1× PBS (pH 7.4) and twice with 50 mM ammonium bicarbonate. Proteins on the beads were digested by adding 200 ng trypsin/Lys-C mix (Promega) for 16 h at 37° C. The digests were reduced, alkylated, acidified, and desalted using GL-Tip SDB (GL Sciences Inc., Tokyo, Japan), and the eluates were evaporated and dissolved in 0.1% trifluoroacetic acid (TFA) and 3% acetonitrile.

LC-MS/MS analysis of the resulting peptides was performed on an EASY-nLC 1200 UHPLC connected to an Orbitrap Fusion mass spectrometer through a nanoelectrospray ion source (Thermo Fisher Scientific). Peptides were separated on a 75-µm inner diameter × 150 mm C18 reverse phase column (Nikkyo Technos, Tokyo, Japan) with a linear gradient of 4%-32% acetonitrile for 0-100 min followed by an increase to 80% acetonitrile for 100-110 min. The mass spectrometer was operated in a data-dependent acquisition mode with a maximum duty cycle of 3. MS1 spectra were measured with a resolution of 120,000, an automatic gain control (AGC) target of 4e5, and a mass range of 375 to 1,500 *m/z*. Higher-energy collisional dissociation-MS/MS spectra were acquired in the linear ion trap with an AGC target of 1e4, an isolation window of 1.6 *m/z*, a maximum injection time of 35 ms, and a normalized collision energy of 30. Dynamic exclusion was set to 20 s. Raw data were analyzed directly against the Swiss-Prot database restricted to *Mus musculus* using Proteome Discoverer version 2.4 (Thermo Fisher Scientific) for identification and label-free precursor ion quantification. Search parameters were as follows: (a) trypsin as an enzyme with up to two missed cleavages; (b) precursor mass tolerance of 10 ppm; (c) fragment mass tolerance of 0.6 Da; and (d) cysteine carbamidomethylation as a fixed modification; and (e) protein N-terminal acetylation and methionine oxidation as variable modifications. Peptides were filtered with a false discovery rate of 1% using the percolator node. Normalization was performed so that the total sum of abundance values for each sample was equal across all peptides.

##### Neo-amino-terminal MS

To determine candidate OMA1 substrates during mitochondrial stress, cells were treated with or without 40 μM FCCP for 2 h, then washed once with 1× PBS (pH 7.4), and cell pellets were collected. The pellets were then lysed in a guanidine buffer (6 M guanidine-HCl, 100 mM HEPES-NaOH [pH 7.5], 10 mM TCEP [Sigma Aldrich], and 40 mM 2-chloroacetamide [CAA, Sigma Aldrich]). After heating and sonication, 30 μg of proteins was purified by methanol-chloroform precipitation and resuspended in 20 μL of PTS buffer (100 mM Tris-HCl [pH 8.0], 12 mM sodium deoxycholate, and 12 mM sodium lauroylsarcosinate). The protein solution was diluted 10-fold with 10 mM CaCl_2_ and digested with 600 ng of Tryp-N (LysargiNase, Merck Millipore) overnight at 37° C. After acidification with 0.5% TFA (final conc.), an equal volume of ethyl acetate was added to each sample, followed by centrifugation at 15,700*g* for 2 min to separate the ethyl acetate layer. The aqueous layer was collected and desalted using GL-Tip SDB, and the eluates were evaporated and dissolved in 50 μL of 2.5% formic acid and 30% acetonitrile. Enrichment of protein N-terminal peptides was performed using GL-Tip SCX (GL Sciences Inc.) based on the previous report (Chang *et al*, 2021). Flow-through fractions were evaporated and dissolved in 0.1% TFA and 3% acetonitrile. LC-MS/MS analysis of the resulting peptides was performed as described above. Raw data were analyzed directly against the Swiss-Prot database restricted to *Mus musculus* using Proteome Discoverer version 2.4 for identification and label-free precursor ion quantification. Search parameters were as follows: (a) Tryp-N as a semi-specific enzyme with up to two missed cleavages; (b) precursor mass tolerance of 10 ppm; (c) fragment mass tolerance of 0.6 Da; (d) cysteine carbamidomethylation as a fixed modification; and (e) protein N-terminal acetylation and methionine oxidation as variable modifications.

##### Activities of respiratory chain complexes

Mitochondrial fractions from each Flp-In T-REx-293 cell line were solubilized in 20 mM Bis-Tris (pH 7.0) buffer containing 2 mM NaCl, 500 mM aminocaproic acid, 10% (w/v) glycerol, 1 mM EDTA, 0.5% (w/v) digitonin, and protease inhibitor cocktail for 15 min on ice, and the extracts were centrifuged at 15,000*g* for 10 min at 4° C. The clarified supernatant was then applied to CN-PAGE, followed by evaluation of in-gel activities by incubating the gel in either Complex I (NADH dehydrogenase) activity substrate (5 mM Tris-HCl [pH 7.4], 0.2 mM NADH, and 0.25% [w/v] nitro blue tetrazolium [NBT] chloride) or Complex II (succinate dehydrogenase) activity substrate (5 mM Tris-HCl [pH 7.4], 20 mM succinate, 0.2 mM phenazine methosulfate, and 0.25% [w/v] NBT). The activities of Complexes I and II were quantified using ImageJ software (Schneider *et al*, 2012).

#### Metabolite measurement

For metabolite extraction, 4 × 10^6^ cells were treated with methanol containing internal standards (H3304-1002, Human Metabolome Technologies, Inc. (HMT), Tsuruoka, Japan), including AMP, ADP, and ATP. The extract was then centrifuged at 2,300*g* for 5 min at 4° C. The resulting supernatant was filtered using a 5-kDa cutoff centrifugal filter at 9,100*g* and 4° C to remove macromolecules. The filtrate was then dried and reconstituted in Milli-Q water for metabolomic analysis. Anion analysis was performed by HMT using capillary electrophoresis- (CE-) -MS/MS, as previously described (Ohashi *et al*, 2008; Ooga *et al*, 2011). CE-MS/MS analysis was performed using an Agilent CE capillary electrophoresis system equipped with an Agilent 6460 Triple Quadrupole LC/MS (Agilent Technologies, Inc., CA). Peak areas of individual metabolites were calculated using MasterHands software (Keio University, Tsuruoka, Japan) (Sugimoto *et al*, 2010) and MassHunter Quantitative Analysis (Agilent). Peak areas of AMP, ADP, and ATP were normalized to their respective internal standards, and concentrations were determined using standard curves.

#### Measurement of cellular respiration

The oxygen consumption rate was measured in a Seahorse Extracellular Flux Analyzer XFe96 (Agilent) in cells grown in DMEM-GlutaMAX containing 25 mM glucose medium. In each well, 5 × 10^4^ cells were plated and incubated at 37° C for 1 h prior to measurement. ATP production was assessed after the addition of oligomycin (2 μM), maximal respiration after FCCP (0.5 μM), and nonmitochondrial respiration after the addition of rotenone and antimycin A (each 0.5 μM).

#### Measurement of ΔΨ_m_

To measure the ΔΨ_m_, 6 × 10^4^ cells were seeded on 96-well plates at 37° C for overnight. The next day, the medium was removed and replaced with fresh medium without (DMSO) or with 20 µM FCCP and incubated for another 1 h. The medium was then replaced with a medium containing 200 nM TMRM (Thermo Fisher Scientific) and incubated for 30 min. Cells were washed once with ice-cold 1× PBS (pH 7.4) and replaced with fresh PBS buffer containing 2% FBS. Fluorescence was measured using an Envision 2105 multimode plate reader (PerkinElmer) at excitation/emission wavelengths of 544 nm/579 nm. Each signal intensity was normalized to the average intensity of DMSO-treated WT cells.

#### Quantification of mtDNA

Total genomic DNA from cultured cells was extracted and purified using the DNeasy Blood and Tissue Kit (Qiagen) and diluted to each concentration of 25 ng/μL. To quantify the amount of mitochondrial DNA (mtDNA) per nuclear DNA (nDNA), quantitative real-time polymerase chain reaction (qPCR) was performed using PowerSYBR Green PCR Master Mix (Thermo Fisher Scientific) and QuantStudio 5 Real-Time PCR System (Thermo Fisher Scientific). Quantification of relative copy number differences was performed based on the difference in the threshold amplification between mtDNA and nDNA [ΔΔC(t) method]. The following primer sets were used for mtDNA: forward (5’-tcgaaaccgcaaacatatca) and reverse (5’-caggcgtttaatggggttta) targeting human ND5; and for nDNA: forward (5’-ctgtggcatccacgaaacta) and reverse (5’-agtacttgcgctcaggagga) targeting human ACTB.

#### Whole cell proteomics

##### Protein digestion

Protein (15 μg) were subjected to tryptic digestion and the samples were reduced (10 mM TCEP) with alkylated (20 mM CAA) in the dark for 45 min at 45° C. Samples were then subjected to an SP3-based digestion (Hughes *et al*, 2014). Washed the SP3 beads (Sera-Mag magnetic carboxylate modified particles, Sigma Aldrich) were mixed equally, and 3 µL of bead slurry was added to each sample. Acetonitrile was added to a final concentration of 50% and washed twice using 70 % ethanol (200 µL) on an in-house manufactured magnet. After an additional acetonitrile wash (200µL), 5 µL digestion solution (10 mM HEPES [pH 8.5] containing 0.5µg Trypsin [Sigma Aldrich] and 0.5µg Lys-C [Wako Pure Chemical Industries]) was added to each sample and incubated overnight at 37° C. Peptides were desalted on a magnet using 2 × 200 µL acetonitrile. Peptides were eluted in 10 µL 5% DMSO in LC-MS water (Sigma Aldrich) in an ultrasonic bath for 10 min. Formic acid and acetonitrile were added to a final concentration of 2.5% and 2%, respectively. Samples were stored at −20° C before subjection to LC-MS/MS analysis.

##### Liquid Chromatography and Mass Spectrometry

LC-MS/MS instrumentation consisted of an Easy-nLC 1200 (Thermo Fisher Scientific) coupled via a nano-electrospray ionization source to an Exploris 480 mass spectrometer (Thermo Fisher Scientific). For peptide separation, an Aurora Frontier column (60 cm × 75 μm C18 UHPLC, IonOpticks, Victoria, Australia) was used. A binary buffer system (A: 0.1 % formic acid and B: 0.1 % formic acid in 80% acetonitrile) based gradient was utilized as follows at a flow rate of 185 nL/min; a linear increase of buffer B from 4% to 28% within 100 min, followed by a linear increase to 40% within 10 min. The buffer B concentration was further increased to 50 % within 4 min, then to 65 % over the next 3 min. The column was then washed with 95 % buffer B for an additional 3 min. The RF Lens amplitude was set to 45%, the capillary temperature was 275° C, and the polarity was set to positive. MS1 profile spectra were acquired using a resolution of 120,000 (at 200 *m*/*z*) at a mass range of 450-850 *m*/*z* and an AGC target of 1 × 10^6^.

For MS/MS independent spectra acquisition, 34 equally spaced windows were acquired at an isolation *m*/*z* range of 7 Th, and the isolation windows overlapped by 1 Th. The fixed first mass was 200 m/z. The isolation center range covered a mass range of 500–740 m/z. Fragmentation spectra were acquired at a resolution of 30,000 at 200 m/z using a maximal injection time setting of ‘auto’ and stepped normalized collision energies (NCE) of 24, 28, and 30. The default charge state was set to 3. The AGC target was set to 3e6 (900% - Exploris 480). MS2 spectra were acquired in centroid mode. FAIMS was enabled using an inner electrode temperature of 100° C and an outer electrode temperature of 90° C. The compensation voltage was set to −45V.

##### Data analysis

Raw files were analyzed using Spectronaut 18.5.231110.55695 (Bruderer *et al*, 2015). In direct DIA (Data independent acquisition) mode using the Uniprot *Homo sapiens* (Human) UP000005640 reviewed only, 20,400 protein sequences). Trypsin/P was selected as the cleavage rule using a specific digest type. The minimal peptide length was set to seven and a total of two missed cleavages were allowed. The peptide spectrum match, peptide, and protein group false discovery rate (FDR) were controlled to 0.01. The mass tolerances were used with default settings (Dynamic,1). The directDIA +(deep) workflow was selected and cross-run normalization was enabled. The protein group file was exported and LFQ intensities (MaxLFQ algorithm; Cox *et al*, 2014) were log2 transformed. Statistically significant different proteins were identified using a two-sided *t*-test followed by a permutation-based FDR calculation (s0=0.1, number permutations=500, FDR< 0.05) using Instant Clue (Nolte *et al*, 2018).

##### Cell growth assay

For the cell growth assay, we used our customized oxidative medium (glucose-free D-MEM [Thermo Fisher Scientific] supplemented with 10% FBS, 2% GlutaMAX, 1% penicillin [100 U/mL]-streptomycin [100 μg/mL], and 10 mM galactose as a carbon source) to cultivate cell cultures. The day before the assay, 4 × 10^5^ cells were seeded on 6-well dish and incubated overnight at 37°C. Every 2 days (total of 8 days), cells were counted in triplicate using the Countess II automated cell counter (Thermo Fisher Scientific).

### In organello import assay

The *In organello* import assay was performed as previously described with minor modifications (Salscheider *et al*, 2022). Radiolabeled precursor proteins were synthesized by a mammalian cell-free protein synthesis system using the rabbit reticulocyte lysate-based TNT SP6 Quick Coupled Transcription/Translation System (Promega) containing [^35^S]-methionine according to the manufacturer’s protocol. Mitochondria (40 μg) isolated from cells as described above were subjected to each import reaction. The protein import assay was initiated by mixing the radiolabeled precursor protein with the mitochondrial fraction at 30° C in the presence or absence of 40 μM FCCP/10 μM oligomycin A. The import reaction was stopped after 8 or 15 min by placing the reactants on ice. The samples were then treated with proteinase K (10 μg/mL) for 15 min to degrade non-imported precursor protein. After protease inactivation by the addition of 1 mM PMSF, mitochondria were washed twice with homogenization buffer and analyzed by SDS-PAGE, and the radiolabeled proteins were detected by autoradiography using Typhoon FLA 9500 (Cytiva, Marlborough, MA).

#### RNA sequencing

RNA samples used for sequencing analysis were quantified using an ND-1000 spectrophotometer (NanoDrop Technologies, Wilmington, DE) and their quality was confirmed using a Tapestation (Agilent). Sequencing libraries were prepared from 200 ng total RNA using the MGIEasy rRNA Depletion Kit and the MGIEasy RNA Directional Library Prep Set (MGI Tech Co., Ltd.) according to the manufacturer’s protocol. The libraries were sequenced on the DNBSEQ-G400 FAST Sequencer (MGI Tech Co., Ltd.) using the 150 nt paired-end strategy. To analyze the sequencing data, all sequencing reads were trimmed for low-quality bases and adapters using Trimmomatic (v.0.38) (Bolger *et al*, 2014). Raw counts for each gene were estimated in each sample using RSEM version 1.3.0 (Li & Dewey, 2011) and Bowtie 2 (Langmead & Salzberg, 2012). We used the edgeR program (Robinson *et al*, 2010) to detect differentially expressed genes (DEGs). Normalized counts per million (CPM) values and fold-changes (logFC) between samples were obtained from the raw gene-level counts. We defined differentially expressed genes as those exhibiting a fold-change of ≥2, either upregulated or downregulated.

#### Quantification and statistical analysis

Statistical analysis was performed using GraphPad Prism 8.4. We considered different cell populations to be biologic replicates; aliquots or repeated measurements of a cell population were considered technical replicates. We obtained at least three independent reproducible results for most key experiments, although we did not perform explicit power calculations. Data are presented as the mean ± SD, and statistical significance was assessed by one-way analysis of variance (ANOVA) followed by Tukey’s multiple comparison or by two-way ANOVA followed by Bonferroni’s multiple comparison test. A *p*-value <0.05 was considered statistically significant. Chi-square tests with Bonferroni correction were also used to compare categorical data between groups.

## Data availability

All data contained in the main text and the supporting information are available upon reasonable request. The IP-MS proteomics data have been deposited to the ProteomeXchange Consortium via the jPOST partner repository with the dataset identifiers PXD063014, PXD063017, and PXD063094 (https://repository.jpostdb.org/). The mitochondrial proteome data were deposited to the PRoteomics IDEntifications (PRIDE) database (https://www.ebi.ac.uk/pride/) with the dataset identifier PXD063602. The RNA sequencing data were deposited to the Gene Expression Omnibus (GEO) with the dataset identifier GSE296062 (https://www.ncbi.nlm.nih.gov/geo/).

## Expanded View

This article contains supporting information.

## Acknowledgements

We thank David Chan (Caltech) and Naotada Ishihara (Osaka University, Japan) for kindly providing the *OPA1*^−/−^ and *Drp-1*^−/−^ MEFs, respectively. The *OMA1*^−/−^ MEFs were kindly provided by Carlos López-Otín (University of Oviedo, Spain), and the plasmid pcDNA3.1(+)/Su9-TEV was gifted from Shiori Sekine (University of Pittsburgh). We also thank Takashi Tatsuta (Max Planck Institute) for fruitful discussions on the study, Kohei Nishino (Tokushima University, Japan) for assistance with the proteomic analysis, Robin Alexander Rothemann (University of Cologne, Germany) for generating the AIFM1 cell line, Seitaro Koga for critical suggestions on the Western blotting analysis, and Takahiro Yoshinaka for early work on this study. This work was supported by the JSPS KAKENHI Grants (No. 25K14949 to M.N. and 21K19389, 23K26789, and 25H02336 to T.K.), the Deutsche Forschungsgemeinschaft (DFG, German Research Foundation) - SFB 1218 – Projektnummer 269925409 (to T.L. and J.R.), Medical Research Center Initiative for High Depth Omics (to H.K.), and the Joint Usage and Joint Research Programs, the Institute of Advanced Medical Sciences, Tokushima University (to T.K.).

## Author contributions

**Mitsuhiro Nishigori:** Formal analysis, data curation, validation, investigation, visualization, methodology. **Serina Hirata**: Data curation, formal analysis, investigation, methodology. **Hidetaka Kosako:** Data curation, formal analysis, methodology. **Hendrik Nolte:** Data curation, formal analysis, investigation, methodology. **Jan Riemer:** Data curation, methodology. **Thomas Langer:** Conceptualization, funding acquisition, supervision, writing – review & editing.

**Takumi Koshiba:** Conceptualization, data curation, funding acquisition, supervision, writing – original draft.

In addition to the CRediT author contributions listed above, the contributions in detail are: T.K., T.L., and M.N. designed most of the experiments, and M.N. and S.H. performed most of the planned biochemical and cell culture experiments. H.K. and H.N. performed the LC-MS/MS experiments and analyzed the proteomic data. J.R. established the *AIFM1* KO in the Flp-In T-REx-293 cell system and contributed to the *In organello* import assay. T.K. and T.L. supervised the overall direction, managed the funding acquisition, and validated the study by analyzing all the data. T.K. contributed to writing the original draft and T.L revised and edited the manuscript. All the authors analyzed the data, interpreted all the experimental results, and provided comments and edits on the manuscript.

