## Supplementary Table and Figures for "OMA1-mediated cleavage of AIFM1 upon mitochondrial stress and suppression of cell growth through the control of OXPHOS activity"

**File including**

Table EV1. Neo-amino-terminal identified by mass spectrometry
Figure EV1. Biochemical validation of stably expressed OMA1/Myc
Figure EV2. Stress-inducible AIFM1 processing by OMA1
Figure EV3. AIFM1 cleavage site targeted by OMA1 and its non-cleavable variant
Figure EV4. Scheme for generating an N-terminal specific antibody against AIFM1
Figure EV5. Role of OPA1 in OMA1-dependent AIFM1 processing
Figure EV6. Characterization of AIFM1$^{TCS}$ and AIFM1$^{TCS/TEV}$

#### Table EV1. Neo-amino-terminal identified by mass spectrometry

| Gene Symbol | Protein Name | Annotated Sequence |
|---|---|---|
| AIFM1 | Apoptosis-inducing factor 1, mitochondrial | [R].VMGLGLSPEE.[K] |
| AK2 | Isoform 2 of Adenylate kinase 2, mitochondrial | [R].GIHCAIDASQTPDIVFASILAAFSKATS.[-]<br>[R].QAEMLDDLME.[K] |
| ALDH1L2 | Mitochondrial 10-formyltetrahydrofolate dehydrogenase | [R].TPQPEEGATYEGIQ.[K] |
| ALDH4A1 | Delta-1-pyrroline-5-carboxylate dehydrogenase, mitochondrial | [K].STGSVVGQQPFGGA.[R] |
| BCS1L | Mitochondrial chaperone BCS1 | [L].SVAPQQSLVLLEDVDAAFLS.[R] |
| CRLS1 | Cardiolipin synthase (CMP-forming) | [K].AAPEPAAGGGGAAAQAPSA.[R] |
| DNAJA3 | DnaJ homolog subfamily A member 3, mitochondrial | [K].QYDAYGSAGFDPGTSSSGQGYW.[R] |
| GLUD1 | Glutamate dehydrogenase 1, mitochondrial | [R].DDGSWEVIEGY.[R] |
| GOT2 | Aspartate aminotransferase, mitochondrial | [A].SSWWTHVEMGPPDPILGVTEAFKRDTNS.[K] |
| GPT2 | Alanine aminotransferase 2 | [K].LLEETGICVVPGSGFGQ.[R] |
| HSPD1 | 60 kDa heat shock protein, mitochondrial | [I].AEDVDGEALSTLVLN.[R]<br>[K].CEFQDAYVLLSE.[K] |
| LAP3 | cytosol aminopeptidase | [L].MESPANEMTPT.[R] |
| MRPL46 | 39S ribosomal protein L46, mitochondrial | [K].ALTPLQEEMAGLLQQIEVE.[R]<br>[W].MLPQVEWQPGETL.[R] |
| MRPS7 | 28S ribosomal protein S7, mitochondrial | [K].AAAATETSSVFADPVIS.[K] |
| NNT | NAD(P) transhydrogenase, mitochondrial | [R].EANSIVITPGYGLCAA.[K] |
| PCK2 | Phosphoenolpyruvate carboxykinase [GTP], mitochondrial | [R].QCPIMDPAWEAPEGVPIDAIIFGG.[R] |
| PRDX5 | Peroxiredoxin-5, mitochondrial | [K].ATDLLLDDSLVSLFGN.[R] |
| SFXN3 | Sideroflexin-3 | [R].AGVATPGLTEDQLW.[R] |
| STOML2 | Stomatin-like protein 2, mitochondrial | [K].ESMQMQVEAE.[R] |
| SUCLG1 | Succinate--CoA ligase [ADP/GDP-forming] subunit alpha, mitochondrial | [F].AAAAINEAIDAEIPLVVCITEGIPQQDMV.[R]<br>[Q].SAGVVVSMSPAQLGTTIY.[K] |
| TIMM50 | Mitochondrial import inner membrane translocase subunit TIM50 | [M].IIEPTSPCLLPDPL.[R] |
| TOMM40 | Mitochondrial import receptor subunit TOM40 homolog | [A].SSPPAGPPPPPTPSLVGLPPPPPSPPGFTLPPLGGGLGTGSSTG.[R] |
| UQCRC1 | Cytochrome b-c1 complex subunit 1, mitochondrial | [L].QSVPETQVSILDNGL.[R] |

Fig. EV1

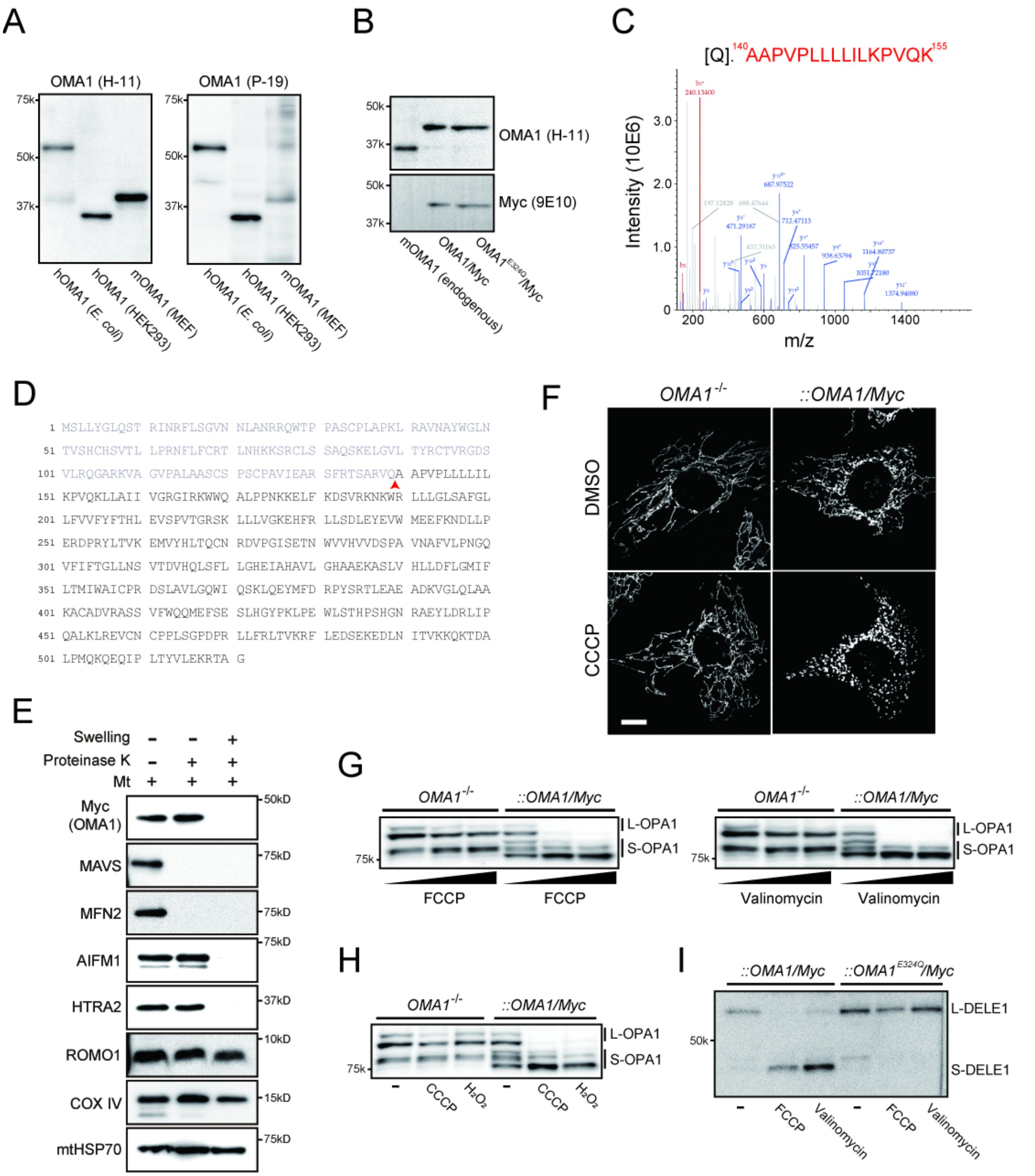


**Figure EV1. Biochemical validation of stably expressed OMA1/Myc.**

**A** Constitutive maturation of OMA1 in mitochondria. The molecular size of OMA1 expressed bacterially (hOMA1) or endogenously in HEK293 (hOMA1) or MEF (mOMA1) cells was compared by SDS-PAGE and Western blotting analysis using anti-OMA1 antibodies (left blot, H-11; right blot, P-19). Size differences in OMA1 between recombinant and endogenous OMA1 show that it is constitutively converted into mature forms after import into mitochondria.

**B** Western blot analysis of the stably expressed Myc-tagged version of WT OMA1 and its E324Q variant in *OMA1*-null MEFs used in the study. These recombinants show constitutive conversion to their mature form as well as to the endogenous (WT MEFs) protein (left lane). The size difference between the endogenous and recombinant proteins indicates the weight of the Myc tag (4-kDa) fused to the C-terminus of each recombinant protein.

**C, D** The N-terminal sequence of OMA1/Myc was determined by MS. Mitochondrial extract from $OMA1^{-/-}$ MEFs stably expressing OMA1/Myc was immunoprecipitated with anti-Myc antibody, followed by tryptic digestion and LC-MS/MS to identify its N-terminal peptide sequence. The MS/MS spectrum suggested that the most distal N-terminal residue was A140 (red peptide, $^{140}$AAPVPLLLLILKPVQK$^{155}$) identical to that of endogenous OMA1 (Baker *et al*, 2014). Mature OMA1 is generated upon proteolytic cleavage at the site indicated by the red arrowhead in (D).

**E** Submitochondrial localization of OMA1/Myc demonstrating its proper localization in the IMS. Mitochondria isolated from $OMA1^{-/-}$ MEFs stably expressing OMA1/Myc were treated with proteinase K (50 μg/mL) under either isotonic (−SW) or hypotonic swelling (+SW) buffer conditions and kept on ice for 15 min. The reactants were developed by immunoblotting with antibodies against Myc or various mitochondrial membrane markers as indicated. MOM proteins: MAVS and MFN2. IMS proteins: AIFM1 and HTRA2. MIM proteins: ROMO1 and COX IV. Matrix protein: mtHSP70.

**F** Mitochondrial morphologies under normal or depolarized conditions. $OMA1^{-/-}$ or $OMA1^{-/-}$ MEFs stably expressing OMA1/Myc were treated with either DMSO or CCCP, and their mitochondrial morphologies were visualized by immunofluorescence microscopy using mtHSP70 antibody. OMA1/Myc controls mitochondrial dynamics upon mitochondrial stress induced by CCCP. Scale bar, 10 μm.

**G, H** $OMA1^{-/-}$ or $OMA1^{-/-}$ MEFs stably expressing OMA1/Myc were treated for 3 h with (G, left blot) FCCP (0, 20, and 40 μM), (G, right blot) valinomycin (0, 0.1, and 1 μg/mL), or (H) $H_2O_2$ (1 mM) and analyzed by immunoblotting with OPA1 antibody. The proteolytic activity of OMA1/Myc was indistinguishable from that of the endogenous protein, and could be used to monitor OMA1-mediated constitutive OPA1 processing under steady state conditions and stress-inducible OMA1 activations. L- and S-OPA1 indicate long- and short-form OPA1, respectively.

**I** $OMA1^{-/-}$ MEFs stably co-expressing both Myc-tagged versions of OMA1 variants (WT and

E324Q) and HA-tagged DELE1 were treated for 3 h with either FCCP (40 μM) or valinomycin (1 μg/mL) and analyzed by immunoblotting with an HA antibody to monitor DELE1 processing in the cells. OMA1/Myc, but not the proteolytically inactive $OMA1^{E324Q}$/Myc, rescued cells processed DELE1 when mitochondrial stress was induced. L- and S- DELE1 indicate long and short form DELE1, respectively.

### Fig. EV2

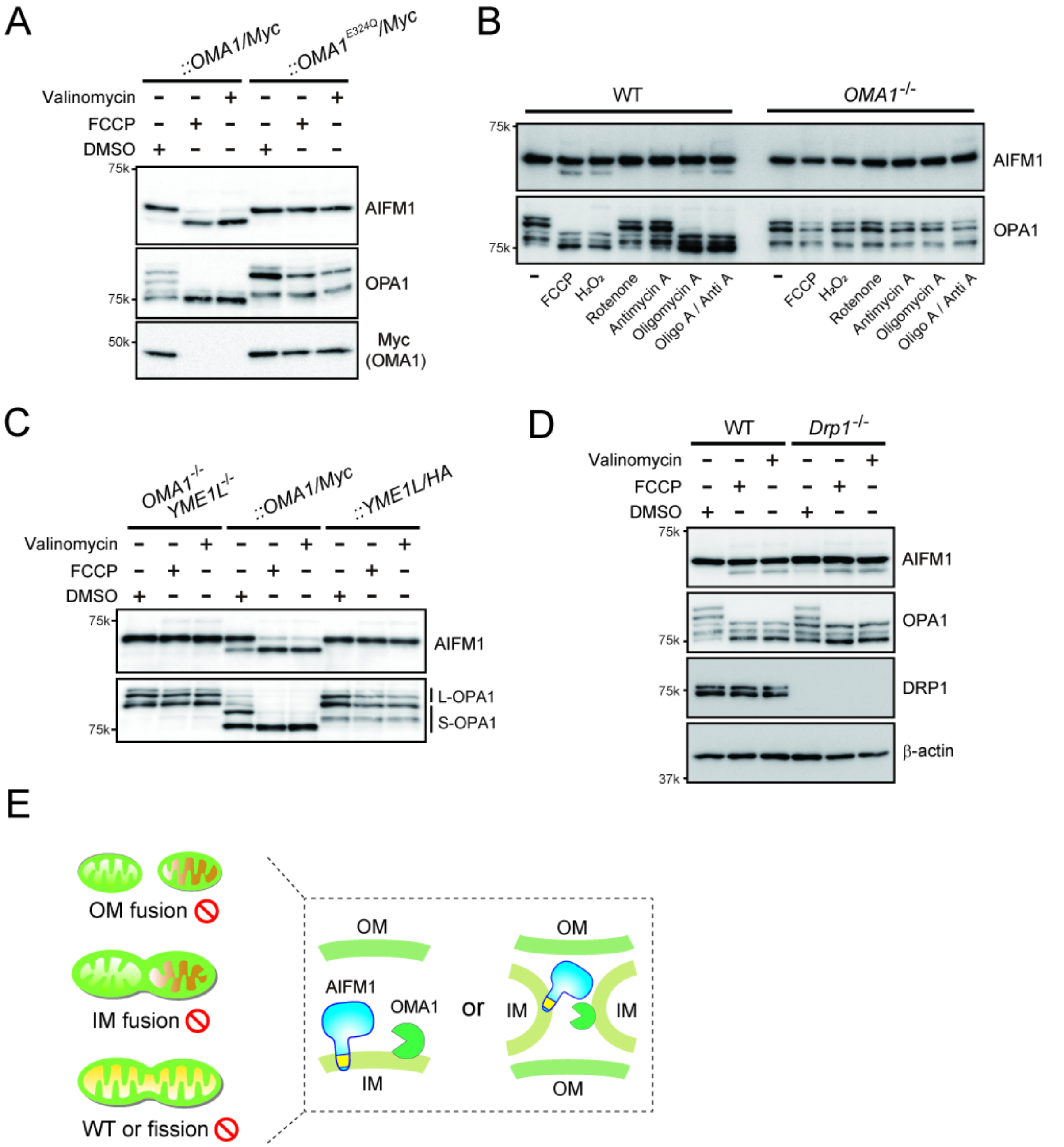


**Figure EV2. Stress-inducible AIFM1 processing by OMA1.**

**A** *OMA1*$^{-/-}$ MEFs stably expressing OMA1/Myc or OMA1$^{E324Q}$ /Myc were treated for 3 h with either FCCP (40 μM) or valinomycin (1 μg/mL) and analyzed by immunoblotting with the indicated antibodies. OMA1$^{E324Q}$ /Myc accumulated stably in cells upon the loss of $\Delta\Psi_m$, whereas OMA1/Myc was rapidly degraded by autocatalytic proteolysis.

**B** Stress-induced substrates (AIFM1 and OPA1) processed by OMA1. *OMA1*$^{+/+}$ (WT) and *OMA1*$^{-/-}$ MEFs were incubated for 3h with FCCP, $H_2O_2$, Rotenone, Antimycin A (Anti A), Oligomycin A (Oligo A), or Anti A/ Oligo A and analyzed by Western blotting.

**C** *OMA1*$^{-/-}$*YME1L*$^{-/-}$ MEFs expressing either OMA1/Myc or YME1L/HA were incubated for 3 h in the absence (DMSO) or presence of either FCCP (40 μM) or valinomycin (1 μg/mL) and analyzed by immunoblotting (indicated antibodies).

**D** OMA1-dependent processing of AIFM1 in *Drp1 KO* cells. The WT or *Drp1* KO MEFs were treated for 3 h with either FCCP (40 μM) or valinomycin (1 μg/mL) and analyzed by immunoblotting with the indicated antibodies.

**E** Illustration shows a hypothetical model of OMA1-mediated AIFM1 processing on the same membrane (acting in *cis*) or on different membranes (in *trans*). *Mfns*-DKO and *OPA1*-KO are incompetent for OM and IM fusion, respectively. *Drp1*-KO is incompetent for OM fission.

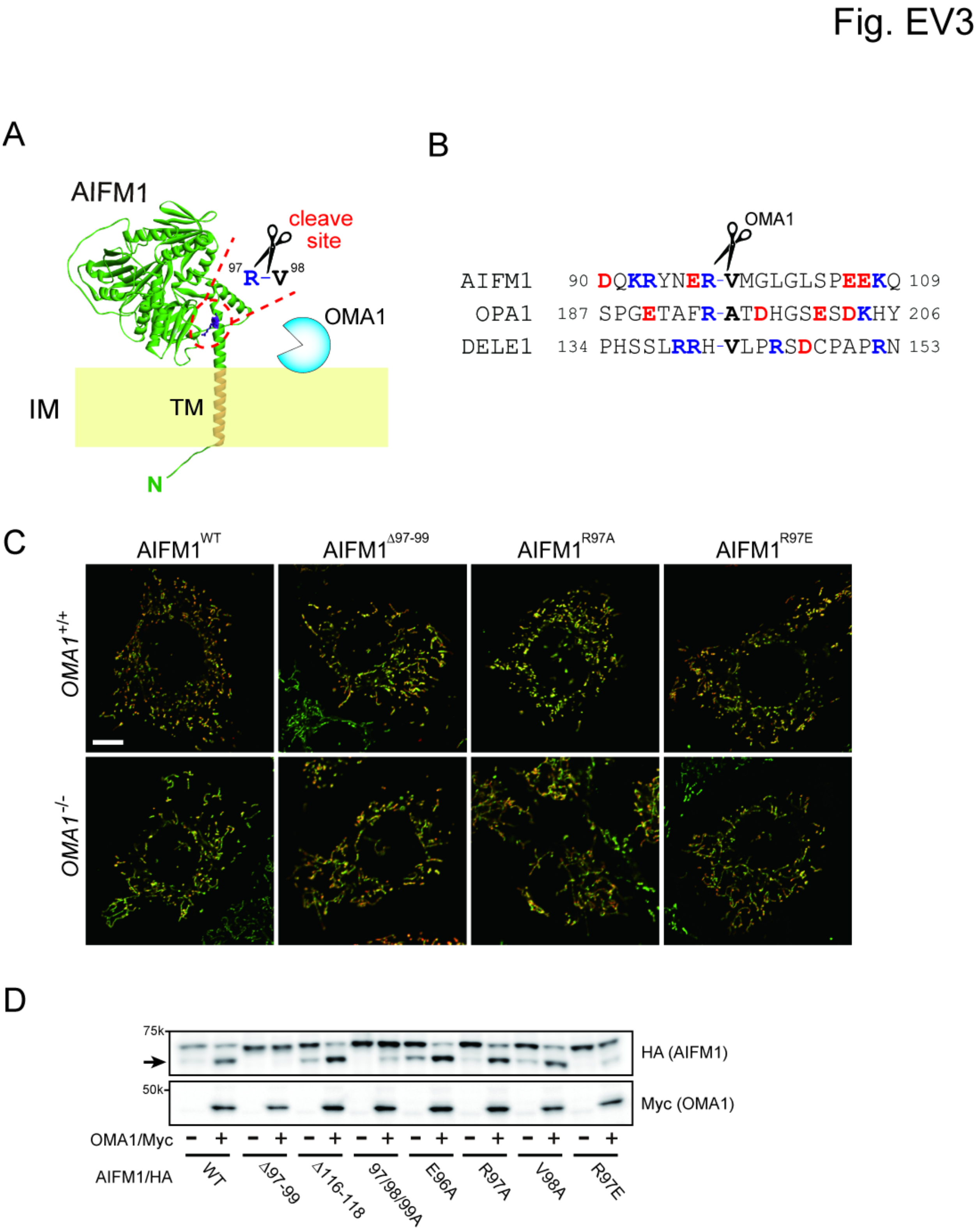


**Figure EV3. AIFM1 cleavage site targeted by OMA1 and its non-cleavable variant.**

**A** Topologic model of the cleavage site of AIFM1 targeted by OMA1. The cleavage site (between residues Arg$^{97}$ and Val$^{98}$) is located just downstream of the transmembrane (TM) domain in MIM-anchored AIFM1 and is depicted as a stick model (purple) in the structure. The structure of the mature form of murine AIFM1 (amino acids 55 to 612) as assigned by the AlphaFold3 program (Jumper *et al*, 2021).

**B** Sequence alignment of OMA1 substrates in *Mus musculus*. No conservation of any OMA1 cleavage motif is observed among these substrates.

**C** The indicated Myc-labeled WT or AIFM1 variants (Δ97-99, R97A, and R97E) were stably expressed in either *OMA1*$^{+/+}$ or *OMA1*$^{-/-}$ MEFs by a retroviral system. Immunofluorescence against the Myc epitope was used to identify AIFM1 variants in cells and to determine their subcellular localization (red). Mitochondria in the same cells were also identified by staining with anti-mtHSP70 antibody (green). Both AIFM1 and mtHSP70 in mitochondria were completely merged (yellow) in all cells observed. Scale bar, 10 μm.

**D** HA-tagged plasmids encoding WT AIFM1 or its variants were co-transfected with the Myc-tagged version of OMA1 into HEK293 cells. OMA1-mediated AIFM1 processing was analyzed by Western blotting with anti-HA monoclonal antibody. The arrow indicates the cleaved AIFM1 band.

Fig. EV4

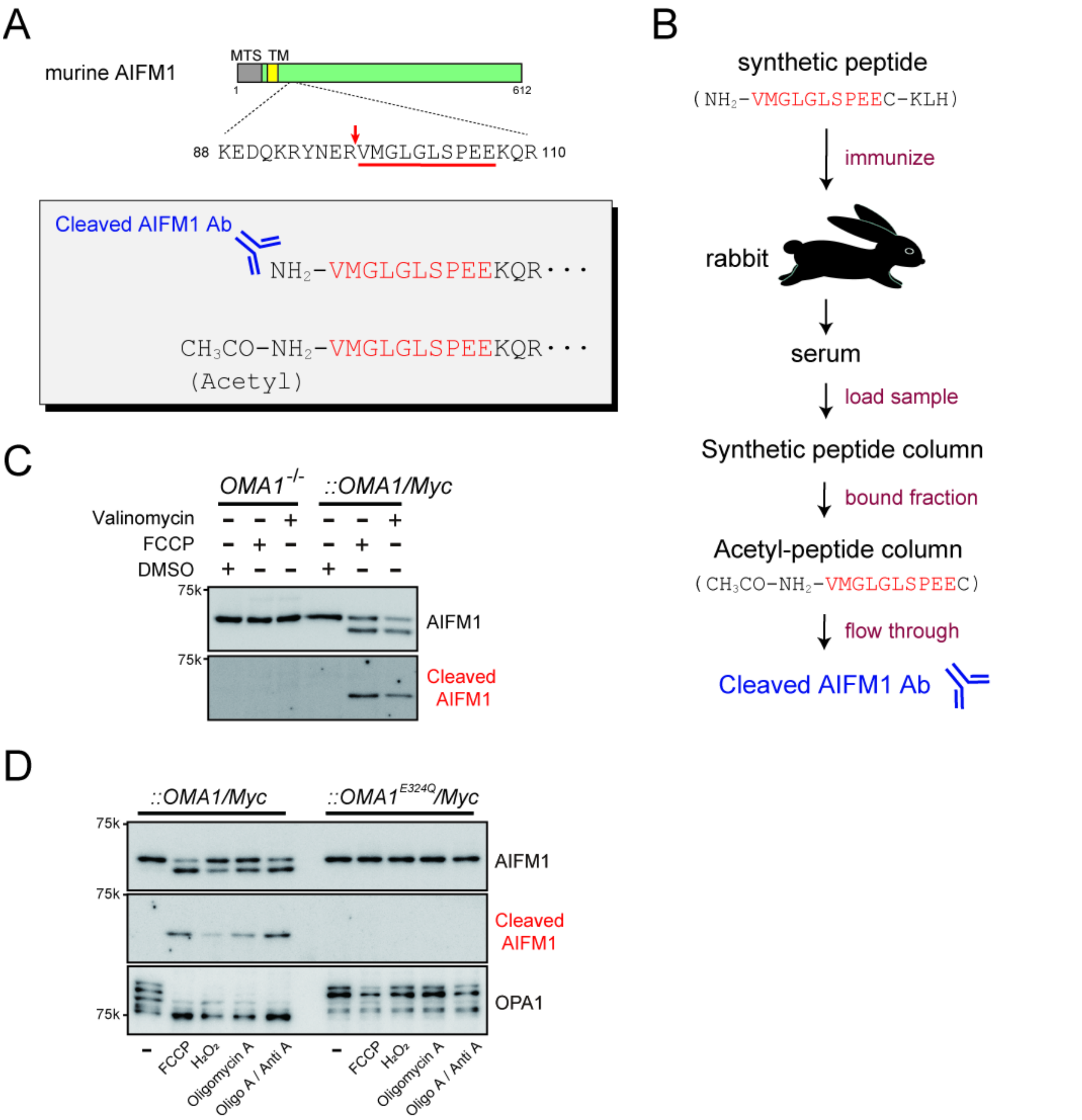


**Figure EV4. Scheme for generating an N-terminal specific antibody against AIFM1.**

**A** Position of the AIFM1 cleavage site (red arrow) targeted by OMA1. The sequence portion underlined in red was chemically synthesized for immunized rabbits to generate polyclonal antibodies against the region (lower inset). The same N-terminally acetylated peptide was also synthesized for use in affinity purification.

**B** Flowchart of the generation of the custom antibody against the N-terminal portion of AIFM1.

**C, D** *OMA1*$^{-/-}$ or *OMA1*$^{-/-}$ MEFs stably expressing OMA1/Myc were treated for 3 h with either FCCP (40 μM) or valinomycin (1 μg/mL) and analyzed by immunoblotting with the custom cleaved AIFM1 antibody (bottom). The antibody specificity was confirmed by comparing the same immunoblot with that detected by the normal AIFM1 antibody (top). In (D), *OMA1*$^{-/-}$ MEFs stably expressing OMA1/Myc or OMA1$^{E324Q}$ /Myc were treated for 3 h with the indicated chemicals and analyzed by immunoblotting with the indicated antibodies.

Fig. EV5

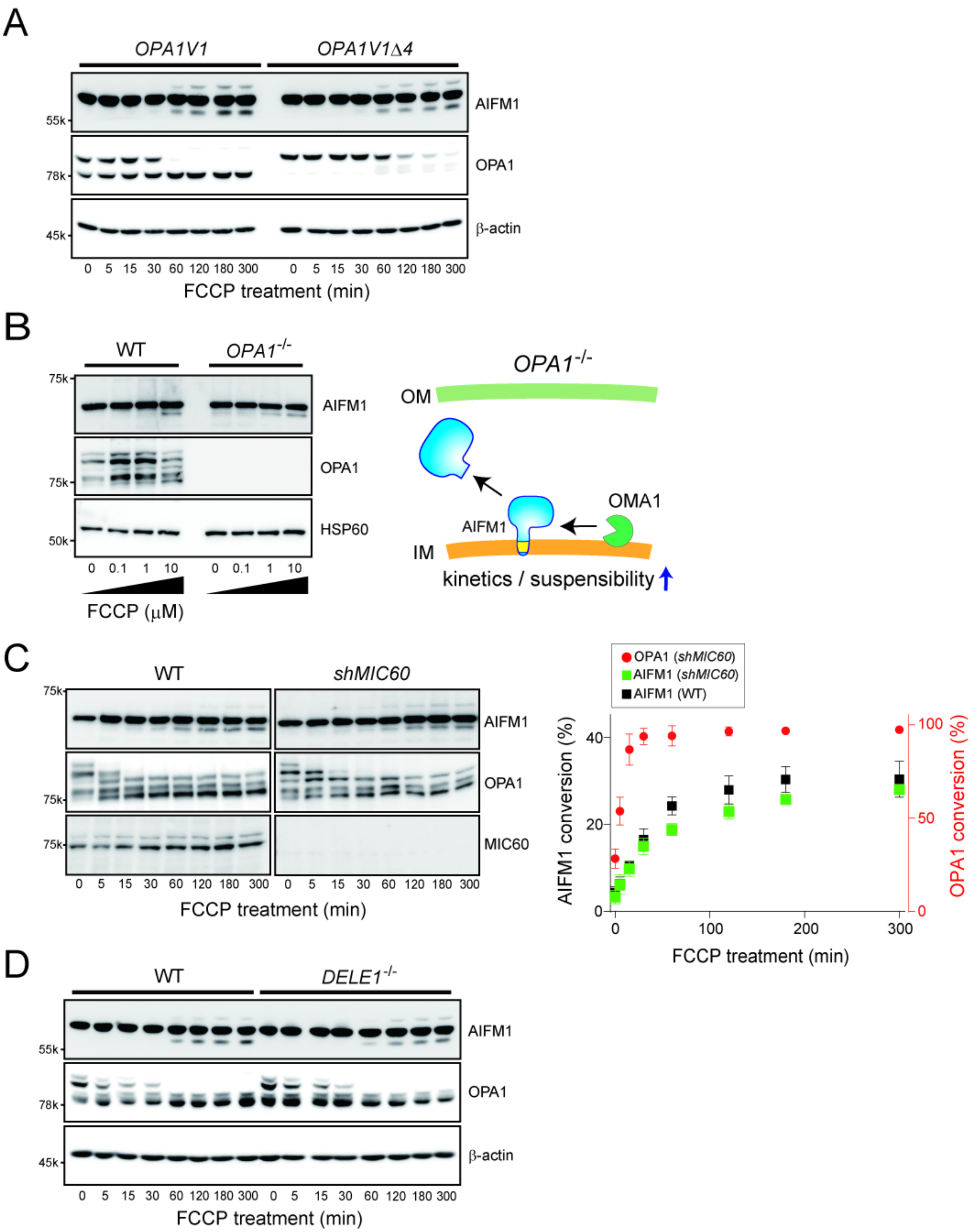
A
OPA1V1
OPA1V1Δ4
AIFM1
OPA1
β-actin
55k
78k
45k
0 5 15 30 60 120 180 300 0 5 15 30 60 120 180 300
FCCP treatment (min)
B
WT
OPA1-/-
75k
AIFM1
OPA1
75k
HSP60
50k
0 0.1 1 10 0 0.1 1 10
FCCP (μM)
OPA1-/-
OM
OMA1
AIFM1
IM
kinetics / suspensibility
C
WT
shMIC60
75k
AIFM1
OPA1
75k
MIC60
75k
0 5 15 30 60 120 180 300 0 5 15 30 60 120 180 300
FCCP treatment (min)
OPA1 (shMIC60)
AIFM1 (shMIC60)
AIFM1 (WT)
AIFM1 conversion (%)
OPA1 conversion (%)
FCCP treatment (min)
D
WT
DELE1-/-
AIFM1
OPA1
β-actin
55k
78k
45k
0 5 15 30 60 120 180 300 0 5 15 30 60 120 180 300
FCCP treatment (min)

**Figure EV5. Role of OPA1 in OMA1-dependent AIFM1 processing.**

**A** MEFs only expressing either the OPA1 V1 isoform (*OPA1V1*) or its non-cleavable version (*OPA1V1Δ4*) (Ahola *et al*, 2024) incubated with FCCP (40 μM) were collected at the indicated time points (0, 5, 15, 30, 60, 120, 180, and 300 min) and analyzed by Western blot. β-Actin blots were used as loading controls for each time point.

**B** WT or *OPA1*$^{-/-}$ MEFs were treated for 3 h with different concentrations of FCCP (0, 0.1, 1 and 10 μM) and analyzed by immunoblotting with the indicated antibodies. Right, Model of OMA1-mediated AIFM1 processing in the absence of OPA1. The loss of OPA1 in cells increases the susceptibility of AIFM1 to OMA1 attack.

**C** MEFs without or with expression of shRNA against *MIC60* incubated with FCCP (40 μM) were collected at the indicated time points (0, 5, 15, 30, 60, 120, 180, and 300 min) and analyzed by Western blot. Right graph, OMA1-dependent substrate processing in *shMIC60* MEFs (red, OPA1; green, AIFM1) was quantified (*n* = 3 biologic replicates) and their conversions (%) were plotted. AIFM1 processing in WT MEFs (control) is shown in black.

**D** Similar to (A), except that the WT or *DELE1*$^{-/-}$ MEFs were treated with FCCP (40 μM) for the indicated times.

Fig. EV6

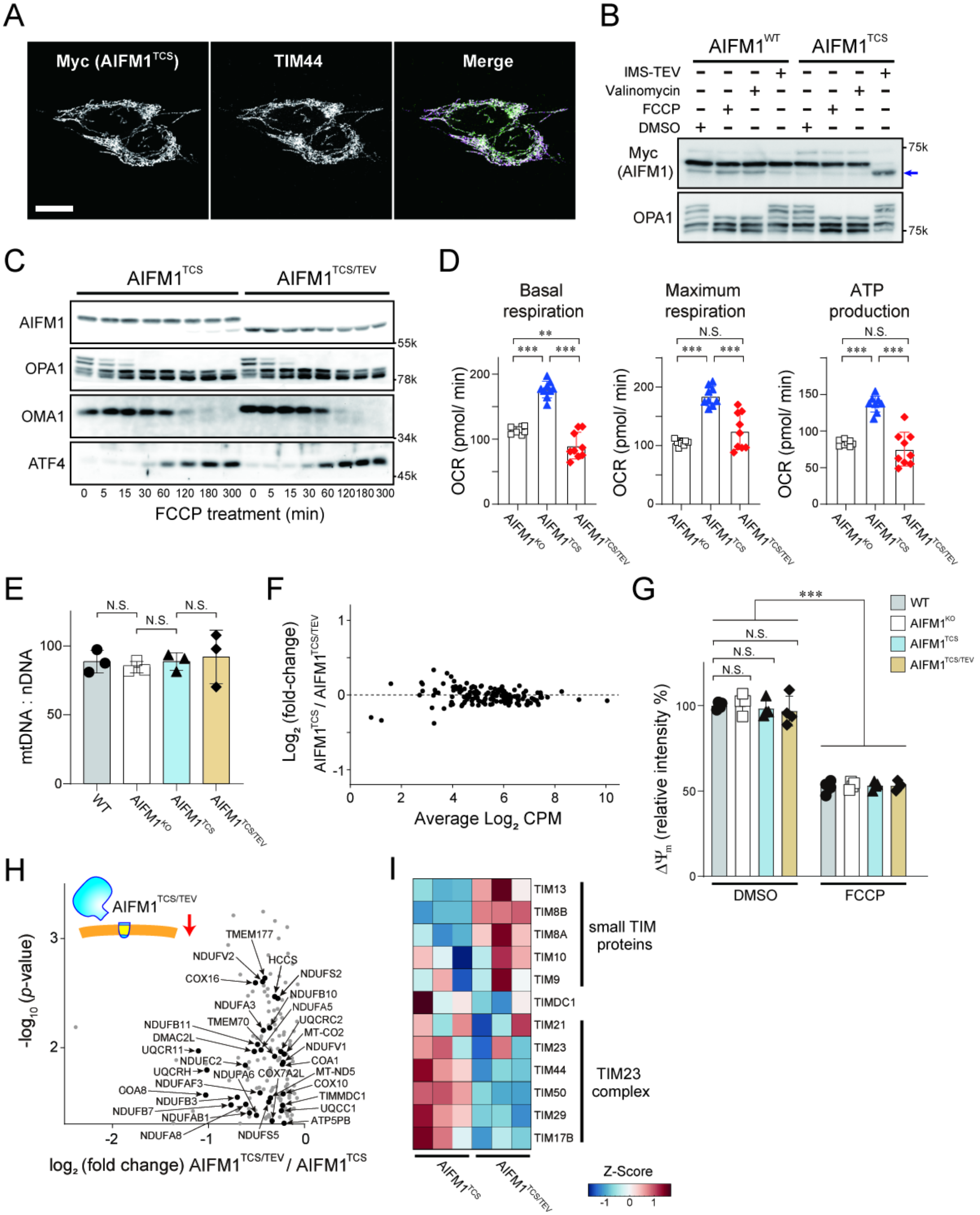


**Figure EV6. Characterization of AIFM1$^{TCS}$ and AIFM1$^{TCS/TEV}$.**

**A** Subcellular localization of AIFM1$^{TCS}$. Flp-In-293-AIFM1$^{TCS}$/Myc cells were monitored by immunofluorescence against the Myc epitope to determine its subcellular localization (left). Mitochondria in the same cells were also identified by staining with an anti-TIM44 antibody (middle). We confirmed that both AIFM1$^{TCS}$ (magenta) and TIM44 (green) were completely merged in the mitochondria (bottom). Scale bar, 10 μm.

**B** The Flp-In-293 AIFM1$^{WT}$/Myc or AIFM1$^{TCS}$/Myc cells were treated for 3 h with either FCCP (40 μM) or valinomycin (1 μg/mL) and analyzed by immunoblotting with the indicated antibodies. AIFM1$^{TCS}$/Myc cells were resistant to the FCCP/valinomycin treatment but able to convert to the cleaved form in the presence of IMS-TEV (right lane). Arrow, cleaved AIFM1.

**C** AIFM1$^{TCS}$- or AIFM1$^{TCS/TEV}$-expressing cells incubated with FCCP (40 μM) were collected at the indicated time points (0, 5, 15, 30, 60, 120, 180, and 300 min) and analyzed by Western blot. In this experiment, we confirmed that OPA1 processing as well as OMA1 and ATF4 activations were similar in both cell types, indicating that the TEV protease targeted to the IMS had no significant off-target effect.

**D** Basal respiration, maximal respiration, and ATP production calculated from the OCR experiment of *AIFM1* KO cells or cells expressing AIFM1 variants (AIFM1$^{TCS}$ and AIFM1$^{TCS/TEV}$) in Fig. 5H. Data shown are mean ± SD ($n$ = 9 biologic replicates). $^{**}p < 0.01$, $^{***}p < 0.001$, and N.S, not significant, respectively.

**E** Analysis of mtDNA copy number per nuclear DNA (nDNA) in WT, *AIFM1* KO, and AIFM1 variant Flp-In-293 cells. Data shown are mean ± SD ($n$ = 3 biologic replicates). N.S., not significant.

**F** Representative MA plot of expressed OXPHOS-related genes (MitoCarta3.0) in AIFM1$^{TCS}$- or AIFM1$^{TCS/TEV}$-expressing cells from transcriptome data. The x-axis represents the $\log_2$ transform of counts per million (CPM), and the y-axis represents the $\log_2$ transform of the fold-change of each gene in AIFM1 variant cells ($n$ = 1). A total of 151 of the OXPHOS-related genes from the transcriptome data were used in this graph. Expression changes greater than or less than 2-fold are considered significant, but we identified no genes meeting these criteria. See also Dataset EV5.

**G** Comparison of the $\Delta\Psi_m$ in WT, *AIFM1* KO, and AIFM1 variant Flp-In-293 cells treated without (DMSO) or with FCCP. Fluorescence values obtained by measuring TMRM were normalized to the average intensity of DMSO-treated WT cells. Data shown are mean ± SD ($n$ = 3 biologic replicates). $^{***}p < 0.001$ and N.S., not significant.

**H** Enlarged box area in Fig. 6A. A total of 123 mitochondrial proteins (MitoCarta3.0) are plotted and 32 of the OXPHOS subunits decreased in AIFM1$^{TCS/TEV}$ cells are labeled. See also Dataset EV4.

**I** Heatmap (Z-scores) showing relative enrichment of selected proteins as identified by IP-MS ($n$ = 3) in Fig. 5A. Components of the TIM23 complex were enriched in the AIFM1$^{TCS}$ precipitates, whereas small TIM proteins showed a higher affinity for the membrane-dislocated AIFM1 variant, $AIFM1^{TCS/TEV}$. See also Dataset EV3.
